# Assessing the performance of the Cell Painting assay across different imaging systems

**DOI:** 10.1101/2023.02.15.528711

**Authors:** Callum Tromans-Coia, Nasim Jamali, Hamdah Shafqat Abbasi, Kenneth A. Giuliano, Mai Hagimoto, Kevin Jan, Erika Kaneko, Stefan Letzsch, Alexander Schreiner, Jonathan Z. Sexton, Mahomi Suzuki, O. Joseph Trask, Mitsunari Yamaguchi, Fumiki Yanagawa, Michael Yang, Anne E. Carpenter, Beth A. Cimini

## Abstract

Quantitative microscopy is a powerful method for performing phenotypic screens from which image-based profiling can extract a wealth of information, termed profiles. These profiles can be used to elucidate the changes in cellular phenotypes across cell populations from different patient samples or following genetic or chemical perturbations. One such image-based profiling method is the Cell Painting assay, which provides morphological insight through the imaging of eight cellular compartments. Here, we examine the performance of the Cell Painting assay across multiple high-throughput microscope systems and find that all are compatible with this assay. Furthermore, we determine independently for each microscope system the best performing settings, providing those who wish to adopt this assay an ideal starting point for their own assays. We also explore the impact of microscopy setting changes in the Cell Painting assay and find that few dramatically reduce the quality of a Cell Painting profile, regardless of the microscope used.

## Introduction

The drug discovery process poses a plethora of challenges, often requiring a specialized combination of solutions. In the early stages of drug discovery, target-based or phenotypic-based approaches can be used to find promising compounds to take forward [1–3]. Phenotypic screens can capture information from vastly complex biological systems and provide a wealth of insight, for example when treating a disease model with a panel of drugs to determine compounds that elicit the desired outcome. The molecular targets of any promising compounds identified through phenotypic screens can then later be elucidated by target deconvolution.

Cell Painting is a phenotypic assay that can be performed at a cost of approximately 26 cents per well for staining reagents at scale in both academia and industry [4,5]. Information-rich data is extracted by imaging eight internal cellular structures using six fluorescent stains, typically over five channels. Across all of these imaged compartments, thousands of features (such as nucleus area, cell shape, staining intensity and more) can be recorded using CellProfiler [6]; alternately, trained deep learning networks can extract features that are more powerful but currently less interpretable [7]. Collections of either type of features are termed profiles [8,9]. These profiles can then be used to elucidate a compound’s mechanism of action by comparing the profile of one compound that has a known mechanism of action with another compound that has an unknown mechanism. Measuring thousands of features at this scale in an unbiased way is known as profiling, which contrasts with screening methods where scientists select individual biologically relevant features of interest. The Cell Painting assay encapsulates a vast wealth of nuanced information about cell state following treatment creating a global view of the cell’s phenotype, rather than a view that is constrained by the preconception of what is expected [10,11]. An approach like this allows virtual drug discovery; for example, the morphological profile of a particular treatment can be queried against other publicly available morphological profiles to reveal perturbations that lead to a similar, or opposing, phenotype [12]. Alternatively, deep learning models can learn to predict assay outcomes for compounds using image-based profiles [13].

In 2019, a collaboration between pharmaceutical companies, non-profit organizations, microscope vendors and several more supporting companies, collectively known as the Joint Undertaking for Morphological Profiling (JUMP) Cell Painting Consortium, was organized. Its goals were to generate a rich trove of imaging data to deepen our understanding of the information that can be derived from Cell Painting images, while also creating a public resource for others to use [5]. The collaborative effort of academic and industry partners in this consortium aimed to reduce the long drug development cycle by making image profiling data open to all. This can in turn drive a new data-driven approach to drug discovery and help to reduce failures of promising compounds in the later stages of the discovery process.

To prepare for the creation of this large public data set, significant efforts were taken to optimize the Cell Painting assay, yielding version 3 of the Cell Painting protocol [14]. Those efforts included the creation of a positive control plate of annotated chemical compounds that would allow benchmarking of Cell Painting assay performance; this plate (JUMP-MOA) contains four replicates each of 40+ pairs of mechanism-of-action-matched compounds, allowing for detection of technical quality in a single 384-well plate [14].

Here, we sought to develop recommendations for settings on multiple high-throughput microscopes from different vendors for the Cell Painting assay. The microscopes tested include those used by different partners in the JUMP Consortium who produced portions of the full dataset at various laboratory sites around the world. Our goal was to optimize the quality of data from a number of different microscopes from different manufacturers for the Consortium, and to be able to confidently recommend Cell Painting imaging settings to a broad variety of end users. Five microscope vendors captured images of the JUMP-MOA standard plates across multiple imaging settings, yielding approximately 450,000 images capturing 41 million cells with a combined size of ∼6.7 TB involving 23 unique microscope setting combinations. We assessed what settings are the most influential and which have a comparatively lower impact in identifying wells treated with the same compound or with drugs that have a similar mechanism of action [15]. Our goal was not to compare or recommend particular instruments; as system configurations can widely vary and end users typically are constrained to use a pre-existing system at their institution. We rather hope to empower users of any microscope system to be able to quickly optimize conditions on whatever microscope they have access to.

## Results and Discussion

We explored the best performing microscope settings combinations to determine best settings for the Cell Painting assay across multiple microscope systems and find that all microscopes tested are fully compatible with the Cell Painting assay. The best performing setting combinations for each microscope are summarized in Tables 1-5. Each “leaderboard” is normalized to the best-performing settings for that microscope and the top scores for all microscopes fall within 6.7% of each other prior to normalization. We used two profile quality metrics: percent replicating (what proportion of compounds correctly match replicates of itself) and percent matching (what proportion of compounds correctly match another compound annotated with the same mechanism of action, MOA), hereafter collectively referred to as profile strength. In tables 1-5, both percent matching and percent replicating are averaged and the best performing setting combination for a given microscope is set to 100% to create the metric called percent score. We additionally sought to tease out any common effects that particular settings have. However, as each vendor independently determined the parameters they wished to test, in many cases we could not conclusively study the effects of individual settings in isolation. Nevertheless, we examined several imaging variables, recognizing that the available data usually did not test each setting independently.

**Table 1:** Molecular Devices ImageXpress Micro Confocal setting performance. The best performing settings within a vendor are set to 100% in the Percent Score column. This normalization was performed individually for each score type. Percent Score is the mean of Percent Replicating and Percent Matching for each individual profile prior to normalization. Duplicate setting combination scores are mean aggregated.

| Place | Modality | Binning | Magnification | NA | Number Of Channels | Z Planes | Sites | Percent Replicating | Percent Matching | Percent Score |
| --- | --- | --- | --- | --- | --- | --- | --- | --- | --- | --- |
| 1 | Widefield | 1 | 20 | 0.75 | 6 | 1 | 9 | 100 | 100 | 100 |
| 2 | Confocal | 1 | 10 | 0.45 | 6 | 3 | 4 | 98.4 | 100 | 98.8 |
| 3 | Widefield | 1 | 10 | 0.45 | 6 | 1 | 4 | 88.5 | 100 | 91.5 |
| 4 | Confocal | 1 | 10 | 0.45 | 6 | 1 | 4 | 91.8 | 80 | 88.8 |
| 5 | Confocal | 1 | 20 | 0.75 | 6 | 1 | 9 | 86.9 | 90 | 87.7 |
| 6 | Confocal | 1 | 20 | 0.75 | 6 | 3 | 4 | 85.2 | 80 | 83.9 |

**Table 2:** Nikon Eclipse Ti2 inverted microscope setting performance. See Table 1 for description.

| Place | Modality | Binning | Magnification | NA | Number Of Channels | Z Planes | Immersion | Sites | Percent Replicating | Percent Matching | Percent Score |
| --- | --- | --- | --- | --- | --- | --- | --- | --- | --- | --- | --- |
| 1 | Widefield | 1 | 20 | 0.75 | 4 | 1 | dry | 9 | 100 | 100 | 100 |
| 2 | Widefield | 1 | 10 | 0.45 | 4 | 1 | dry | 1 | 79.1 | 85.3 | 80.7 |

**Table 3:** Revvity Opera Phenix Plus setting performance. See Table 1 for description.

| Place | Modality | Binning | Magnification | NA | Number Of Channels | Z Planes | Immersion | Sites | Percent Replicating | Percent Matching | Percent Score |
| --- | --- | --- | --- | --- | --- | --- | --- | --- | --- | --- | --- |
| 1 | Confocal | 2 | 20 | 1 | 5 | 1 | water | 3 | 96.8 | 100 | 100 |
| 2 | Widefield | 1 | 20 | 1 | 5 | 1 | water | 3 | 97.2 | 97.5 | 99.6 |
| 3 | Confocal | 2 | 20 | 1 | 5 | 3 | water | 3 | 100 | 87.5 | 98.8 |
| 4 | Widefield | 2 | 20 | 1 | 5 | 1 | water | 3 | 95.9 | 97.5 | 98.6 |
| 5 | Widefield | 2 | 20 | 1 | 5 | 3 | water | 3 | 97.2 | 92.5 | 98.2 |
| 6 | Widefield | 1 | 20 | 1 | 5 | 3 | water | 3 | 97.2 | 90 | 97.5 |
| 7 | Confocal | 1 | 20 | 1 | 5 | 3 | water | 3 | 95.9 | 92.5 | 97.2 |
| 8 | Confocal | 1 | 20 | 1 | 5 | 1 | water | 3 | 94.5 | 90 | 95.5 |

**Table 4:**
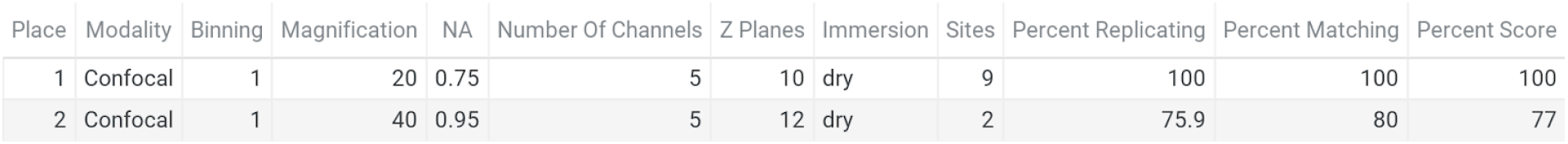
Yokogawa CQ1 setting performance. See Table 1 for description.

| Place | Modality | Binning | Magnification | NA | Number Of Channels | Z Planes | Immersion | Sites | Percent Replicating | Percent Matching | Percent Score |
| --- | --- | --- | --- | --- | --- | --- | --- | --- | --- | --- | --- |
| 1 | Confocal | 1 | 20 | 0.75 | 5 | 10 | dry | 9 | 100 | 100 | 100 |
| 2 | Confocal | 1 | 40 | 0.95 | 5 | 12 | dry | 2 | 75.9 | 80 | 77 |

**Table 5:** Yokogawa CV8000 setting performance. See Table 1 for description.

| Place | Modality | Binning | Magnification | NA | Number Of Channels | Simultaneous Excitation | Z Planes | Immersion | Sites | Percent Replicating | Percent Matching | Percent Score |
| --- | --- | --- | --- | --- | --- | --- | --- | --- | --- | --- | --- | --- |
| 1 | Confocal | 1 | 20 | 1 | 5 | 2 | 12 | water | 9 | 100 | 100 | 100 |
| 2 | Confocal | 1 | 20 | 1 | 6 | 4 | 12 | water | 9 | 91.7 | 100 | 93.8 |
| 3 | Confocal | 1 | 20 | 1 | 5 | 2 | 1 | water | 9 | 90 | 95 | 91.3 |
| 4 | Confocal | 1 | 10 | 0.4 | 6 | 2 | 12 | dry | 4 | 85 | 100 | 88.9 |
| 5 | Confocal | 1 | 20 | 1 | 6 | 1 | 12 | water | 9 | 88.3 | 80 | 86.2 |
| 6 | Confocal | 1 | 40 | 1 | 6 | 4 | 12 | water | 9 | 81.7 | 80 | 81.2 |

Researchers must typically evaluate a tradeoff when selecting an objective for a new imaging assay. While higher magnification objectives, especially at higher numerical apertures (NA), can capture more detail, they cover a smaller field of view and thus require more acquisition sites (and thus longer imaging time) to image the same number of cells when compared to lower magnification objectives. We initially found that data sets taken with 20X magnification objectives typically yielded better profile strength when compared to data sets taken with 10X or 40X objectives (Figure 1A). We also found that experiments that imaged three or more sites per well tended to have an increased profile strength (Figure 1B, Figure S1BC). The objective NA did not appear to yield a difference in our sample (Figure S1A). We also observe that percent matching for individual plates often has the same percent matching value as other plates (Figure 1AB). This behavior is due to there being only 43 MoA classes with more than one member that can be compared to find a match, thus limiting the total number of combinations that can be made when compared to the 90 replicates in the case of the percent replicating metric. Additionally, we find that percent matching scores are in a low 16-26% range. Finding MoA matches from image features alone is a remarkably difficult task [12]. A contributor to making this task challenging can be drug polypharmacology, leading to the induction of different cellular morphologies despite drugs being reported to have the same MoA. In contrast, percent replicating achieves much higher scores due to cells being treated with identical compounds. We also see that alternative metrics, such as mean average precision, report somewhat higher values, while being well correlated with the main metrics used here (Figure S2AB).

**Figure 1:**
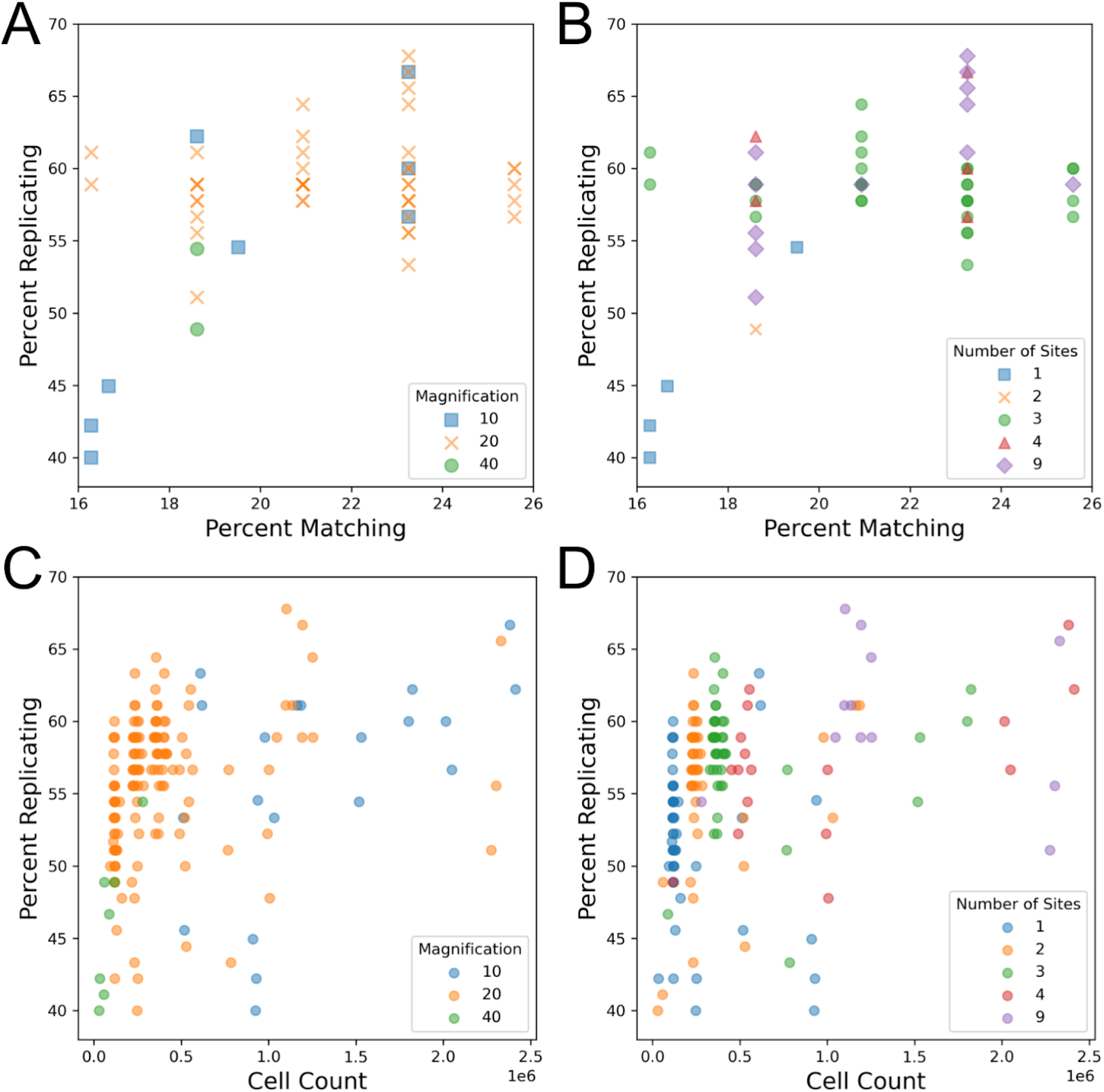
Impact of magnification, number of imaged sites, and cell count on technical quality of the Cell Painting assay. Each data point represents one 384-well plate. Note that the data points are the same in A + B and in C + D but the color labeling changes in each. (A) Comparison of objective magnification in the Cell Painting assay. Because all plates contain the same finite number of replicates and matches, some data points match exactly. (B) Profile strength increases with the number of sites taken per well. (C) An increase in cell count typically leads to an increase in profile strength, leveling off around one million cells. Plot includes artificially site-subsampled data. Cell count shown is sum across all wells. (D) An increase in cell count and profile strength is associated with an increase in the number of sites taken per well. Cell count shown is sum across all wells.

Previous publications [14] and internal data (Way, Singh, and Carpenter, personal communication) have shown that increasing the number of cells per well improves the number of detectable phenotypes. While we observe a positive relationship between site count and profile strength, which could be indicative of taking an improved “sample” of a well (ie. more sites, which can better reveal variations present within a population of cells, such well-edge and center related effects), it could also be a result of imaging more cells. Indeed, in line with previous results we do find that imaging more cells typically leads to a higher profile strength whether measured by percent replicating (Figure 1CD, Figure S1D) or percent matching (Figure S1E).

In order to uncouple the existing inverse relationship present between magnification and cell count, we artificially subsampled profiles that contained more than one acquisition site to all pre-existing site counts (1, 2, 3, 4), creating profiles that had site counts lower than their original (Figure 1CD). While this subsampling does not entirely remove the association between cell count, it allows us to simulate performance across a wider range of cell counts for each magnification than were present in the original data. Here, we see that profile strength increases in tandem with cell count until approximately one million cells per plate (approximately 2,500 cells per well) where we see a mild leveling off. We also show that this effect holds for individual plate profiles: when subsampling the 49 profiles with at least 3 sites per well, 46 show a positive Pearson correlation coefficient between the number of cells in the profile and percent replicating (median: 0.848)(Figure S3AB). These results are consistent with a prior study with controlled variables that found profile strength eventually levels off as a greater sample of a well is taken [14]. This likely explains the low profile strength for 40X as this magnification does not reach a cell count high enough to observe profile leveling-off. While there may be a difference between 10X and 20X in the 0.5 to 1 million cell count range, we cannot be certain based on the data examined here. We also explored the impact objective immersion has on profile strength and find that water seems to perform better than air, but this setting is difficult to detangle from other settings, such as the magnification used (Figure S4A). These results show that, to the degree measurable here, that having sufficient cell count is more important than the details of the objective chosen for overall ability to detect phenotypic signatures. Thus, researchers can feel free to choose the settings that are optimal based on other possible considerations (imaging time, ability to match to existing data, detection of a particular organelle phenotype, etc). We therefore recommend aiming to acquire ∼2,500 cells per well.

We next sought to explore the impact of imaging modalities (confocal vs widefield microscopy) as well as detector binning on profile strength. To control for vendor-to-vendor differences, this was examined for a single vendor who had adjusted both settings within their data sets. Another vendor who tested confocal vs widefield but not binning is shown in Figure S4B. While binning combines adjacent pixels, leading to a decrease in the effective pixel size and thus resolution, we find that increased binning has little impact on profile strength in the contexts of confocal and widefield microscopy (Figure 2AB) using a 20X magnification 1.0NA objective. This indicates that there appears to be no loss in ability to discern overall phenotypic signatures between an effective pixel size of approximately 0.3 vs 0.6 μm. This indicates binning may be especially attractive in particularly large screens due to its typically shorter image acquisition times, reduced file sizes (up to 4-fold for increasing detector binning from 1 to 2), and faster image processing. We also compared detector binning using the alternative metric mean average precision for replicates and MoA matches and found no significant difference between the performance of these two settings (Figure S5AB).

**Figure 2:**
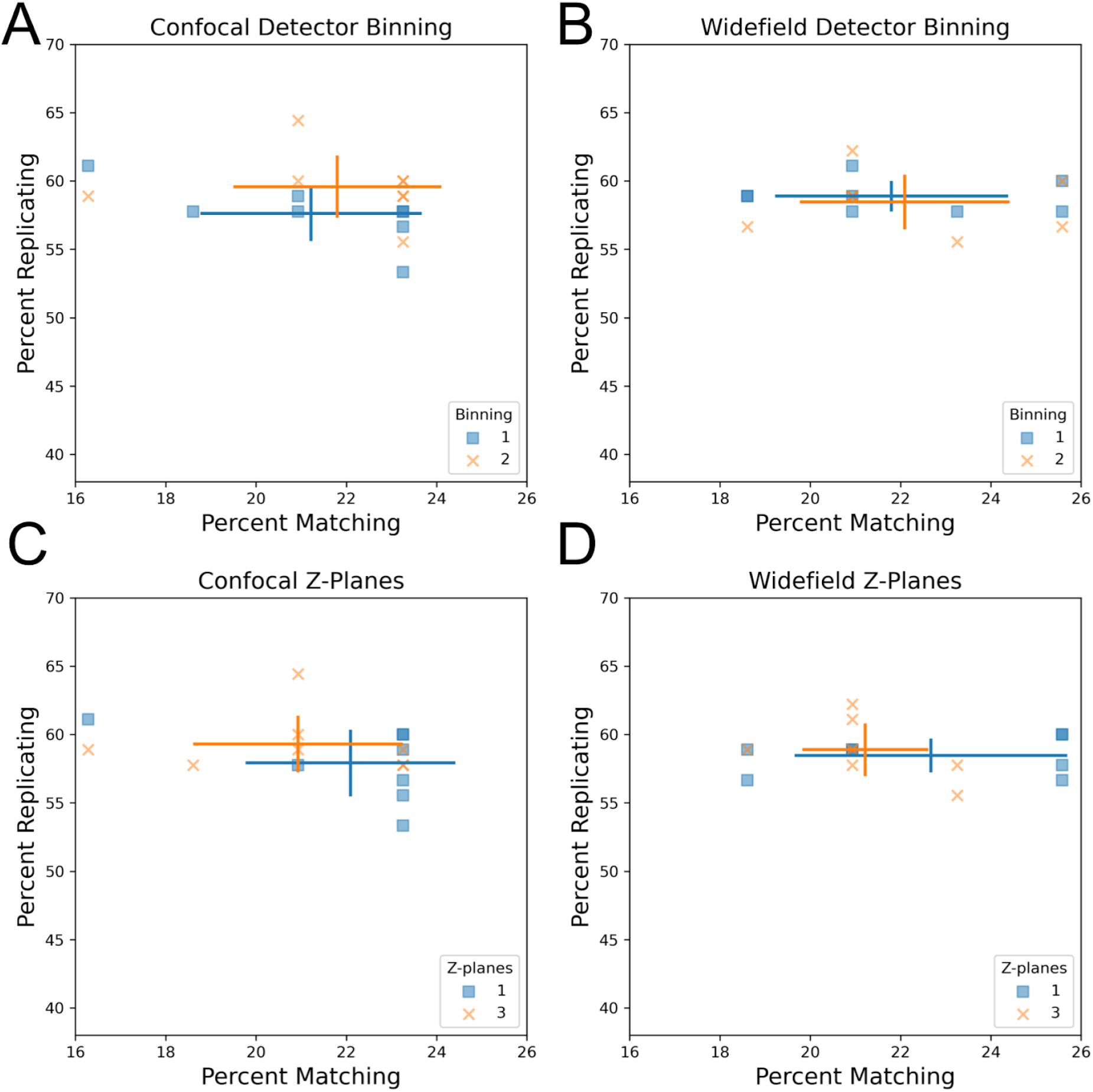
Influence of detector binning and number of z-planes in confocal and widefield microscopy. (AB) Increased detector binning from 1 to 2 has minimal impact on profile strength. Comparison of detector binning in confocal (A) and widefield (B) microscopy. Both widefield and confocal microscopes used the same 20X NA 1.0 lens. Error bars: SD, center of cross: mean for the indicated group. (CD) Increasing the number of max-projected z-planes in confocal and widefield microscopy from 1 to 3 has little impact on profile strength.

We next examined the influence of the number of z-planes on profile strength by comparing single-plane acquisition to multiplane acquisition followed by maximum projection. We found that capturing three z-planes in confocal and widefield microscopy has minimal impact on profile strength compared to a single plane, suggesting that this may be an unnecessary complexity added to Cell Painting imaging workflows (Figure 2CD). However, it is important to note that the U2OS cells imaged here have a flat morphology, so other less flat cell lines may benefit from additional z-planes. We additionally confirmed using mean average precision as an alternative metric that there is no statistically significant difference between one and three z-planes in finding replicate perturbations or MoA matching perturbations (Figure S5CD). Collectively, these results reveal that the computational cost of image processing, storage and acquisition time can be optimized for the Cell Painting assay by increasing detector binning to 2 and reducing the number of z-planes acquired to a single plane.

We next sought to characterize spectral filter effects on image-based profiles. Typically, the Cell Painting assay involves adding six dyes to cells but performing only five-channel imaging due to the difficulty of spectrally separating two of the dye conjugates (Phalloidin/Alexa Fluor 568, which binds to actin, and Wheat-germ agglutinin/Alexa Fluor 555, which binds to the Golgi apparatus and the plasma membrane). This creates the final five effective channels of DNA, endoplasmic reticulum (ER), nucleoli + cytoplasmic RNA (RNA), actin + Golgi + plasma membrane (AGP), and mitochondria (Mito)[4]. Two of these channels (ER and RNA) still share significant spectral similarity but can often be spatially resolved. While the majority of profiles tested used a five-channel acquisition, some omitted taking a separate acquisition of the RNA channel for four total channels and some used alternate filter sets (see Table 6) to separate the AGP channel into actin and Golgi + plasma membrane channels. Unsurprisingly given their close spectral similarity, we find the separated actin and Golgi + plasma membrane channels correlated more strongly with one another than with other channels (Figure S6). We find that six-channel profiles (in which Actin is attempted to be spectrally separated from the Golgi and plasma membrane) did not have any apparent advantage over five-channel profiles (Figure 3A), and therefore find little benefit in attempting this separation unless the researcher is particularly interested in aspects of actin and Golgi biology and wants to study individual phenotypic measurements of these dyes.

**Table 6:**
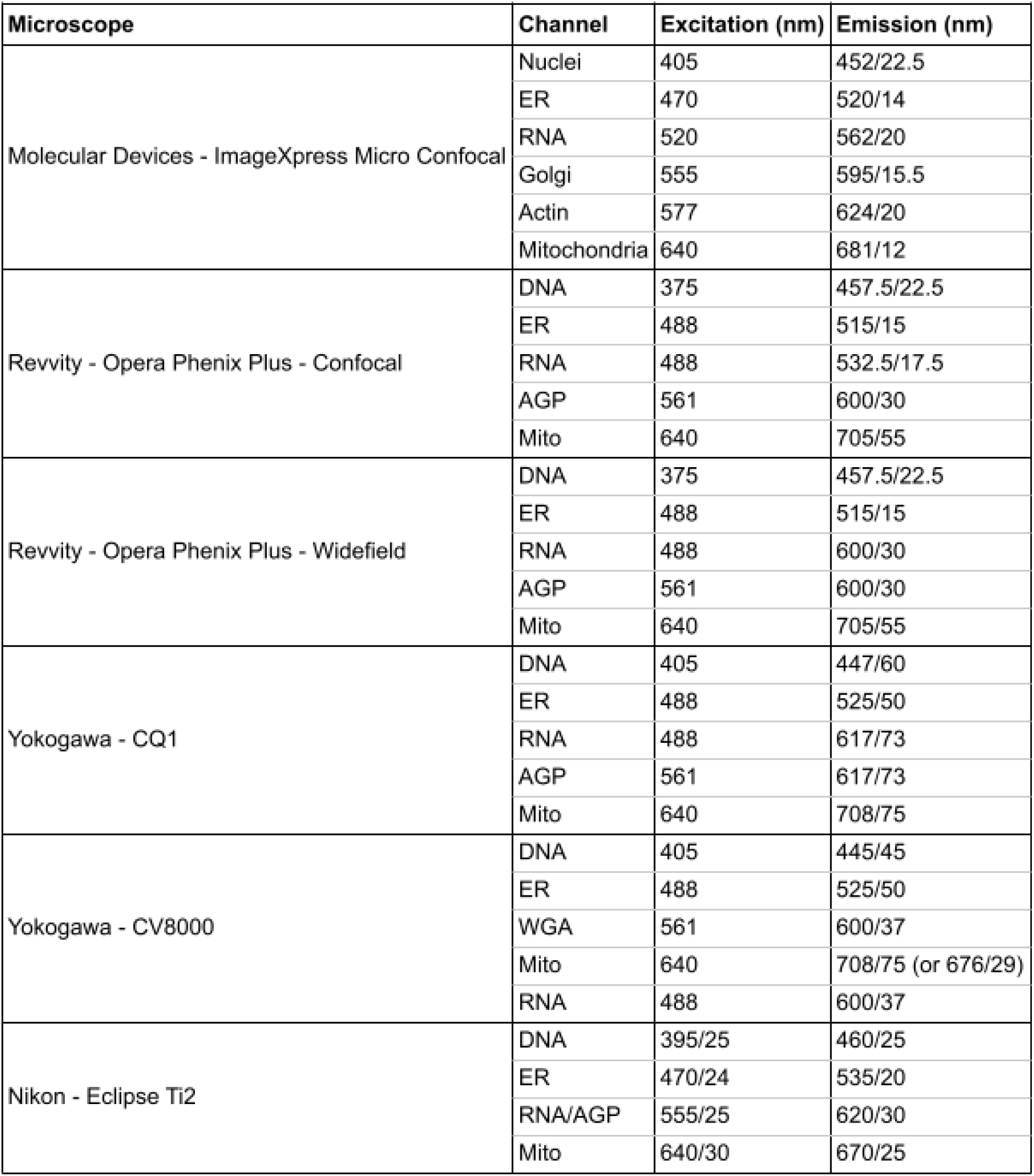
Excitation and emission information for all microscopes tested.

| Microscope | Channel | Excitation (nm) | Emission (nm) |
| --- | --- | --- | --- |
| Molecular Devices - ImageXpress Micro Confocal | Nuclei | 405 | 452/22.5 |
|  | ER | 470 | 520/14 |
|  | RNA | 520 | 562/20 |
|  | Golgi | 555 | 595/15.5 |
|  | Actin | 577 | 624/20 |
|  | Mitochondria | 640 | 681/12 |
| Revvity - Opera Phenix Plus - Confocal | DNA | 375 | 457.5/22.5 |
|  | ER | 488 | 515/15 |
|  | RNA | 488 | 532.5/17.5 |
|  | AGP | 561 | 600/30 |
|  | Mito | 640 | 705/55 |
| Revvity - Opera Phenix Plus - Widefield | DNA | 375 | 457.5/22.5 |
|  | ER | 488 | 515/15 |
|  | RNA | 488 | 600/30 |
|  | AGP | 561 | 600/30 |
|  | Mito | 640 | 705/55 |
| Yokogawa - CQ1 | DNA | 405 | 447/60 |
|  | ER | 488 | 525/50 |
|  | RNA | 488 | 617/73 |
|  | AGP | 561 | 617/73 |
|  | Mito | 640 | 708/75 |
| Yokogawa - CV8000 | DNA | 405 | 445/45 |
|  | ER | 488 | 525/50 |
|  | WGA | 561 | 600/37 |
|  | Mito | 640 | 708/75 (or 676/29) |
|  | RNA | 488 | 600/37 |
| Nikon - Eclipse Ti2 | DNA | 395/25 | 460/25 |
|  | ER | 470/24 | 535/20 |
|  | RNA/AGP | 555/25 | 620/30 |
|  | Mito | 640/30 | 670/25 |

**Figure 3:**
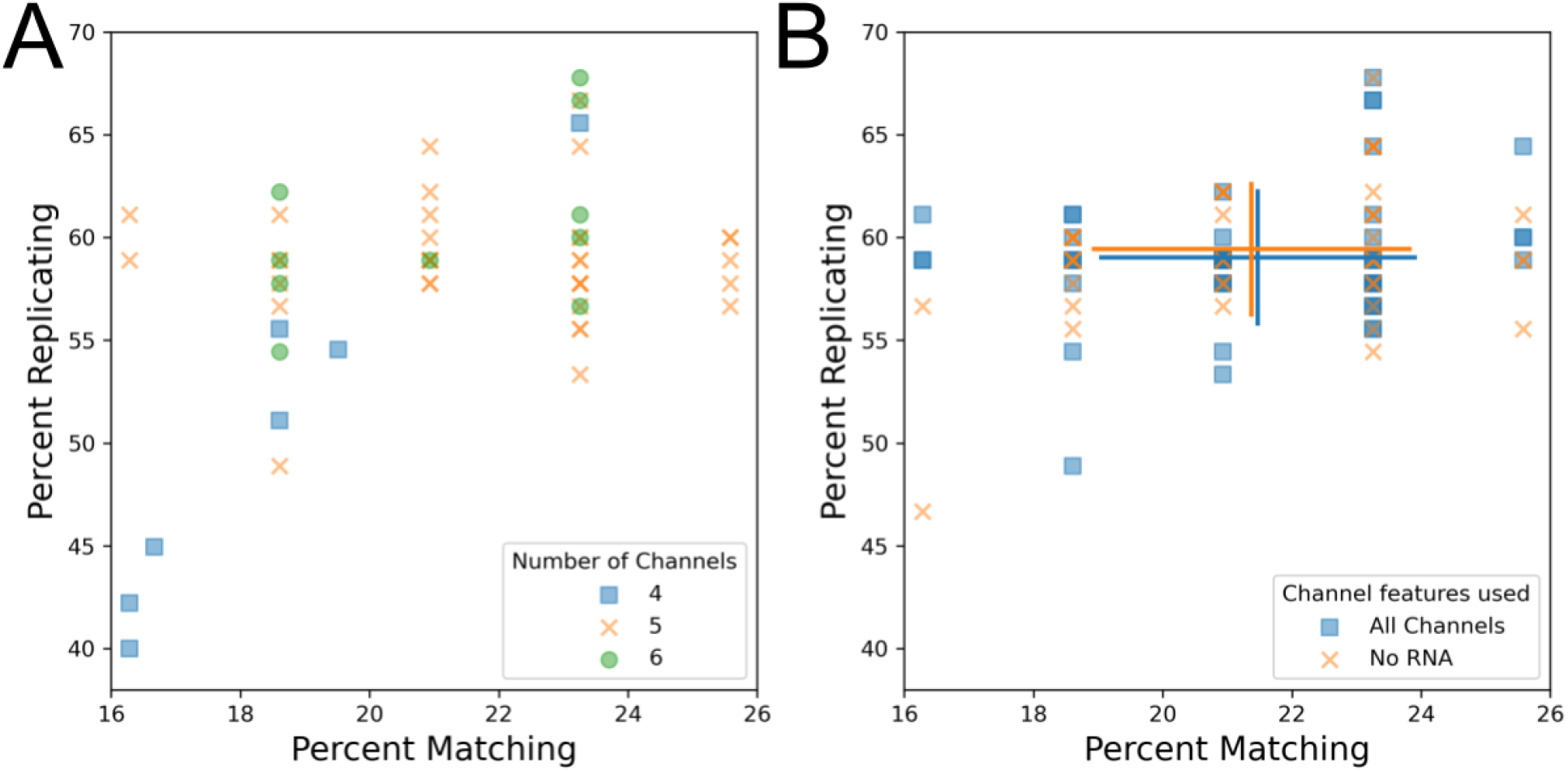
Exploring the technical impact of different channel setups for the Cell Painting assay. (A) Comparison of four, five, and six channel imaging. Since all plates contain the same finite number of replicates and matches, some data points may match exactly. (B) RNA features were dropped prior to feature selection and calculation of percent matching/replicating. Blue squares represent profiles prior to RNA feature dropout and orange Xs represent the same profiles following RNA feature dropout. Error bars: SD, center of cross: mean for a given group.

We found that some profiles from four-channel acquisitions had reduced profile strength (Figure 3A). This result was surprising, given that the dye from the dropped channel (RNA) is typically visible in an alternate channel (ER) and that previous results showed minimal effect of dropping measurement of any channel from the Cell Painting panel [14]. To resolve this discrepancy, we artificially dropped measurements of the RNA channel from profiles which had originally imaged RNA. Consistent with our prior results from a single microscope, we find that artificially dropping RNA features, and thus reducing the number of channels in these profiles, does not noticeably impact profile strength across a sample containing profiles taken on multiple microscopes with multiple spectral setups (88.7% and 89% normalized and mean aggregated percent scores for +RNA and -RNA, respectively) (Figure 3B). We also evaluated the performance of finding replicates and MoA matches using mean average precision as an alternative metric for RNA channel dropout and found that replicate retrieval is significantly impacted by RNA dropout but MoA matching is not (Figure S7AB). This suggests that the reduction in profile strength for the poorly performing aforementioned four-channel profiles could in part be due to a lack of distinct RNA channel but further investigation is needed. Alternatively, since these profiles come from a single site per well, the low performance of the 4-channel profiles could be due to insufficiently high cell count.

One vendor explored using simultaneous excitation, which involves using a system with multiple cameras to capture Cell Painting fluorescence, a process that reduces imaging time. Two different simultaneous excitation setups were tested: two-channel simultaneous excitation was performed on the following channel combinations: endoplasmic reticulum + AGP, mitochondria + RNA, and DNA alone, while four-channel simultaneous excitation was performed on the following channel combinations: DNA + endoplasmic reticulum + AGP + mitochondria, and RNA alone. Filter sets used can be found in Table 6. Profiles generated using four-channel simultaneous excitation minimally impacted profile strength when compared to two-channel, suggesting that simultaneous excitation could be a good additional setting to explore if the imaging system being used supports it (Figure 4A). However, the potential of significant spectral overlap in simultaneous excitation led us to explore if bleedthrough could be observed in the resulting images. Indeed, we find clear bleedthrough of signal between channels, namely DNA signal into the endoplasmic reticulum channel in 4 channel simultaneous excitation images (Figure 4BC, Figure S8). Interestingly, this suggests that overall phenotypic profile separation is relatively insensitive to dye bleedthrough, but this should be interpreted with caution in the absence of evaluation with a larger set of compounds or a more challenging task. Furthermore, an increase in bleedthrough comes with decreased ability to investigate the source of individual feature changes, reducing the accuracy of biological insight that can be derived from the assay.

**Figure 4:**
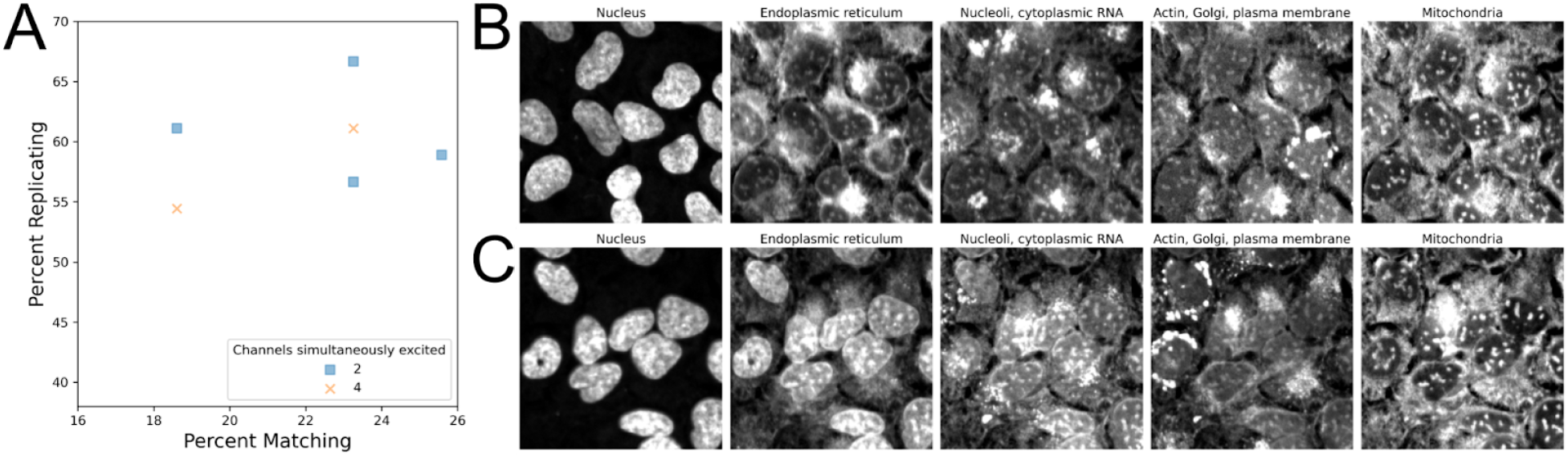
Investigating how simultaneous excitation and spectral overlap can impact the Cell Painting assay. (A) Simultaneous excitation of two channels can be utilized to improve image acquisition times, if available. (BC) Representative images of either two-(B) or four-channel (C) simultaneous excitation. Nuclear signal in (C) can be observed bleeding through into the endoplasmic reticulum channel.

We finally examined how profile strength was impacted by the presence or absence of brightfield images. Unlike the analysis of fluorescent channel z-stacks, which involved maximum projection, brightfield analysis selected a single channel in the middle of the z-stack, corresponding to the plane of best focus. We find that profiles that include brightfield data do not necessarily yield a greater profile strength than those where brightfield data is absent (Figure 5A). We next examined if the profile strength could be improved by removing brightfield features from profiles that originally included this information. We find that dropping the brightfield channel from profiles prior to feature selection has little impact on profile strength (Figure 5B). Profiles with brightfield features had a normalized mean-aggregated percent score of 88% whereas profiles with brightfield features dropped had a score of 92.7%. We also find that when brightfield channel dropping is examined using mean average precision as an alternative metric, there is no statistically significant impact on ability to find MoA replicates or MoA matches (Figure S7CD). While this may suggest that excluding brightfield image acquisition could be considered to improve imaging time, work has previously shown that these images can hold additional insight with more intensive image processing [16]. As these methods may improve in the future, we recommend capturing brightfield for high value public image sets.

**Figure 5:**
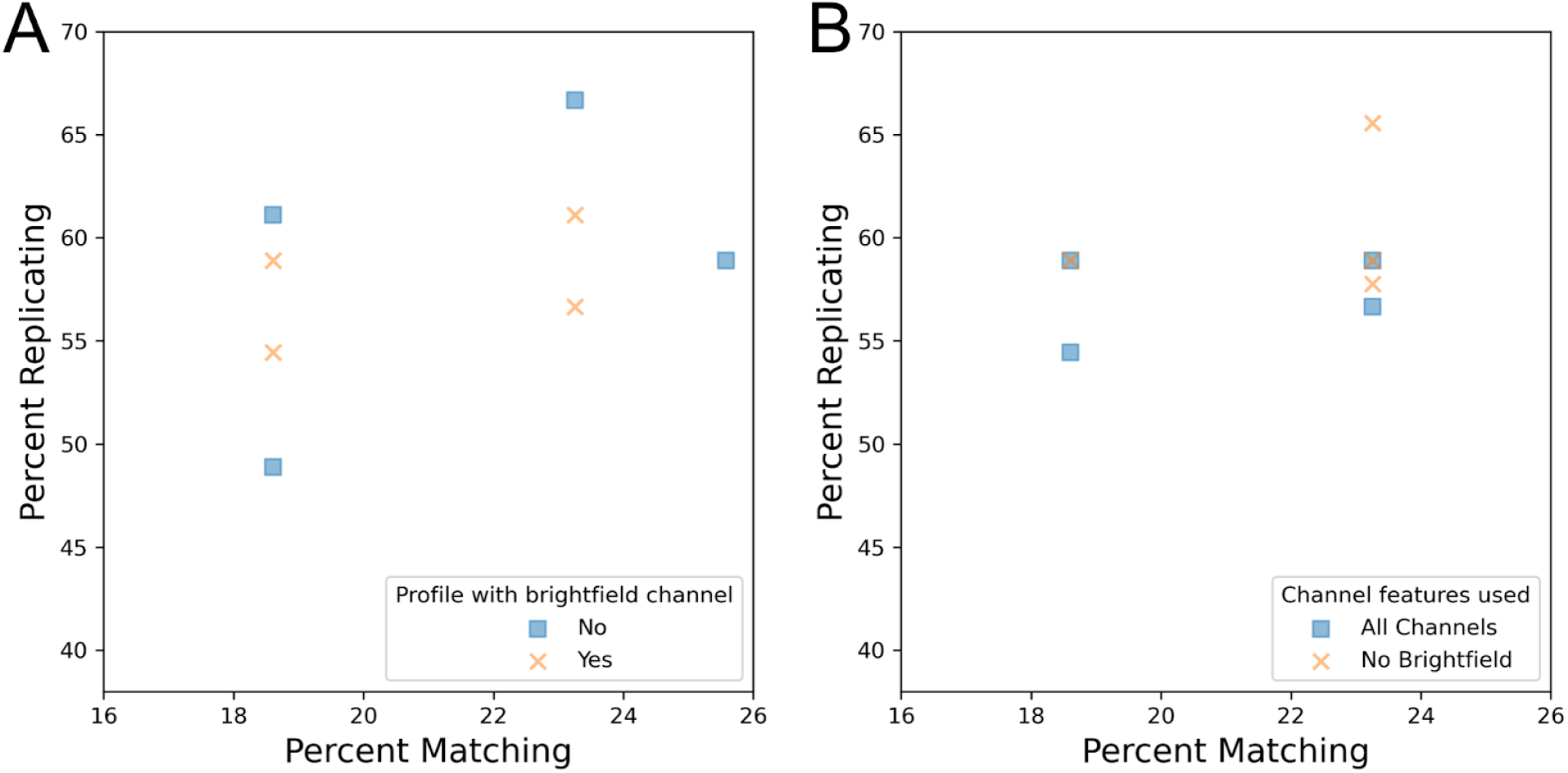
Assessing if acquisition of brightfield images improves the technical quality of the Cell Painting assay. (A) Comparison of profiles that either included or excluded brightfield image acquisition. (B) Brightfield features were dropped from profiles prior to feature selection and calculation of percent replicating/matching.

## Conclusion

In this work we have established the relative performance of various setting configurations across a range of different microscopes for the Cell Painting assay, allowing users to start from an already-optimized set of conditions when beginning to optimize this assay on their own equipment. We hope that these comparisons, as well as our overall recommendations, give confidence to those wishing to adopt the assay, especially as they reveal that there are very few microscopy settings that dramatically decrease the quality of a Cell Painting profile no matter which microscope you happen to use. Deeper exploration into the data also revealed that some settings have a greater impact on the Cell Painting assay than others. We have found that setting choices that increase the cell count, such as decreased magnification and increased number of sites, typically has a positive impact on profile strength. However, due to the interconnected nature of these settings with cell count, it is difficult to clearly conclude those which impact profile strength the most. We also found that simultaneous excitation of four channels leads to bleedthrough in the Cell Painting assay, which can negatively impact interpretation of image-derived features in downstream analysis.

We also found that some settings have a comparatively lower impact on the Cell Painting assay, such as the number of z-planes acquired and detector binning. The number of z-planes can be reduced and detector binning increased with little impact on profile strength, both of which can also afford faster image acquisition. Differences in imaging modality, confocal and widefield, also minimally impact the Cell Painting assay. Additionally, we see no detectable difference when brightfield features are included or removed from profiles, but this does not necessarily suggest that brightfield image acquisition should be excluded from imaging assays completely. Developing work is beginning to reveal that deep learning can provide deeper insight into brightfield images. Recently, it was reported that all five fluorescent channels of a Cell Painting assay can be predicted from the brightfield channel alone [16]. Thus, the brightfield images from an assay could be repurposed at a later date to yield additional insight through label free imaging.

As a final note, while our comparison metrics shown here describe the ability to match compounds and treatments across a broad range of phenotypes, many research applications are not solely about the ability to detect many phenotypes but also the ability to detect one special phenotype of interest. Figure S9A shows examples of images from DMSO and AMG900 treatments which show general reproducibility in these experiments, but also show strong changes in particular individual metrics. If individual metrics or phenotypes are important to a user’s application, care should be taken to ensure the settings chosen allow good signal-to-noise in the user’s preferred feature; features (Figure S10A-E) and feature classes (Figure S11AB) will vary substantially in their sensitivity to changes in imaging parameters. Since Cell Painting’s general profiling ability performs well across a broad range of imaging parameters, users can therefore feel confident that optimizations tailored to their most-preferred specific phenotypes should typically not materially harm their ability to create useful multidimensional Cell Painting profiles.

Taken together, a general set of recommendations for the Cell Painting assay could be: to capture five fluorescent channels and brightfield using a magnification of 20X across four to nine sites or ∼2,500 cells per well, whichever comes first. An increase in binning and a reduction in z-planes can also be explored, depending on the unbinned pixel size and the resolution of any features that are especially interesting to the researcher performing the assay. When following these basic guidelines, Cell Painting can be a powerful tool capable of detecting many phenotypes, across many different kinds of microscopes.

## Methods

### Cell Culture and small molecule treatment

Approximately 2,000 U2OS cells (ATCC cat. no. HTB-96, https://scicrunch.org/resolver/RRID:CVCL_0042) were seeded into each well of a 384-well plate. Cells were then allowed to settle at RT for 1-2 hours to reduce plate effects.

Cells were treated with compounds as found in the JUMP-MOA plate at a final concentration of 3μM. The JUMP-MOA plate layout enables testing of 90 compounds from 47 distinct MOA classes and each compound has 4 replicate wells; see [14] and links within.

### Immunofluorescence

20 μL of mitochondrial staining solution (1.5 μM in cell media which dilutes to a final concentration of 500 nM. MitoTracker Deep Red, Invitrogen M22426) was added to directly each well of the 384-well plate prior to media aspiration to achieve a final well volume of 60 μL. Plates were then incubated in the dark at 37°C for 30 minutes. Cells were then fixed by adding 20 μL of 16% (w/v) methanol-free paraformaldehyde (PFA) to each well, bringing the final volume to 80 μL with a concentration of 4% w/v PFA. Plates were then incubated in the dark at RT for 20 minutes. Wells were then washed with 70 μL of 1x HBSS (Invitrogen, 14065-056) 4 times with the final HBSS wash being aspirated. To each well, 20 μL of staining and permeabilization solution was added (1% BSA, Sigma 05470; 0.1% Triton X100, Sigma T9284; 1 μg/mL Hoechst, Invitrogen H3570; 100 μg/mL concanavalin A/Alexa Fluor 488, Invitrogen C11252; 3 μM SYTO 14, Invitrogen S7576; 1.5 μg/mL WGA/Alexa Fluor 555, Invitrogen W32464; 8.25 nM phalloidin/Alexa Fluor 568, Invitrogen A12380) and incubated in the dark at RT for 30 minutes. Wells were then filled with PBS, plates were then sealed using adhesive foil and shipped to vendor partners at 4°C for image acquisition.

### Image acquisition

Each microscope vendor determined the optimal acquisition settings for their system. Individual setting combinations explored for a particular microscope system are summarized in Tables 1-5. Excitation and emission filter sets used for each microscope can be found in table 6.

### Morphological feature extraction

Morphological features were extracted from images using CellProfiler. First, maximum z-projections were applied to any batches of images that contained more than one z-plane for fluorescent channels. For brightfield images, the middle z-plane of best focus was taken forward for subsequent analysis. Next, an illumination correction function was calculated independently for each channel and applied to all images within a given plate. Nuclei, cytoplasm and cells were segmented from which we then extracted colocalization, granularity, intensity, neighbor, size, shape and texture features.

### Morphological profile generation

Features extracted by CellProfiler were combined into SQLite files using cytominer-database [17] mean aggregated per-well using pycytominer [18]. Next, features were normalized followed by feature selection [19].This process resulted in morphological profiles for each plate that was imaged.

### Data analysis

For each profile, percent replicating and percent matching were calculated. Briefly, these metrics can be determined by first calculating the null distribution from the correlation across features for 10,000 random (non-matching) wells. Then, the feature correlation distribution of replicate wells (percent replicating) or MOA matching wells (percent matching) is calculated. The percentage of the replicate or matching distribution that is above the 95th percentile of the null distribution is the percent replicating or percent matching score, respectively [15].

For calculating mean average precision scores (mAP) we used matric (https://github.com/cytomining/evalzoo/tree/main/matric). We use mAP to assess: (1) replicate retrievability and (2) mechanism of action (MoA) matching. In both scenarios, mAP measures the average similarity within a group of profiles in comparison to their similarities to controls or other perturbations. Similarity calculation involves ranking profiles by cosine distance. For replicate retrievability, mAP indicates how well replicates of a compound can retrieve each other with respect to negative controls. For MoA matching, mAP shows how well compounds annotated with the same MoA can retrieve each other with respect to other compounds. For a more in depth description of mAP, see [5].

Marker selection was performed as described in [20]. In brief, feature selected profiles were loaded into Morpheus (available at https://software.broadinstitute.org/morpheus/) and t-test marker selection was performed between the DMSO and AMG900 treatments. The absolute values for the t-test statistic were sorted and the top 10 highest values are reported.

For feature sensitivity analysis, we performed 2-sample Kolmogorov-Smirnov tests [21] between pairs of plates using unnormalized, mean-aggregated profiles; each individual feature’s 384-well distribution was compared to the other plate’s 384-well distribution, and the p value of the comparison extracted. Since the goal was to compare relative rather than absolute feature sensitivity, no correction for multiple tests was performed. The per-feature p-values were averaged across all pairs of plates within an experimental group, which was either all plates from a particular vendor (Figure S11A), all plates from a particular vendor which had the same pixel size (Figures S10, S11B), or all plates from a particular vendor with the same pixel size and simultaneous excitation settings (Figure S10E). In Figure S10, per-feature measurements as described above were mean aggregated by cellular compartment, measurement module, and the base measurement type; all measurement types were then ranked.

## Funding

This work has been funded in part by the Massachusetts Life Sciences Center Bits to Bytes Capital Call program. Funding for this project was also provided by members of the JUMP Cell Painting consortium in addition to the National Institutes of Health (NIH MIRA R35 GM122547 to AEC). Additional funding was provided to BAC from the Chan Zuckerberg Initiative DAF (2020-225720), an advised fund of the Silicon Valley Community Foundation. These funders had no role in study design, data collection and analysis, decision to publish, or preparation of the manuscript.

Funding was provided by Molecular Devices, Nikon, Revvity, Yokogawa Japan, Yokogawa US to fund creation of the plates and the cost of analysis; these funders had a role in study design and data collection and provided comment on the manuscript but had no role in analysis or decision to publish, nor control over the final manuscript contents.

## Author contributions

AEC and BAC conceived and designed the project. BC, AEC, OJK, FY and MH acquired funding. JZS, OJT, KJ, EK, KAG, MY (Nikon), MS, MY (Yokogawa), SL and AS performed laboratory experiments. HSA and JZS processed image data prior to analysis. HSA and NJ performed image analysis. CTC and NJ generated profiles and performed profile analysis. CTC created the figures and drafted the manuscript, which NJ, AEC and BAC helped revise. Supervision was carried out by BC, AEC, KJ and FY.

## Conflict of Interest

AEC serves as a scientific advisor for Recursion and receives honoraria for occasional talks at pharmaceutical and biotechnology companies. OJT, SL and AS are employees of Revvity, Inc. KJ is an employee of Yokogawa Corporation of America. EK, MH, MS and MY are employees of Yokogawa Electronics Corporation. KAG, MY and FY are employees of Nikon Instruments Inc.

### Acknowledgements

The authors appreciate the more than 100 scientists who have contributed to the organization and scientific direction of the JUMP Cell Painting Consortium. We additionally thank our vendor-partners for creation of the images for this project. We especially thank Alexandr Kalinin, Yu Han, and Shantanu Singh from the Broad Imaging Platform for their advice on evaluation metrics.

Co-first authors CTC and NJ contributed equally to this manuscript, and each has the right to list themselves first in author order on their CVs.

## Data availability

All images, single cell profiles and processed files are freely available in the Cell Painting Gallery https://registry.opendata.aws/cellpainting-gallery/ under cpg0002-jump-scope.

## Code Availability

Jupyter Notebooks used to generate the figures in this paper can be found in the following GitHub repo: https://github.com/jump-cellpainting/jump-scope-analysis

## Notes

### Summary of Updates

Addition of new analysis methods, additional clarification

