## Supplementary figures for "Assessing the performance of the Cell Painting assay across different imaging systems"

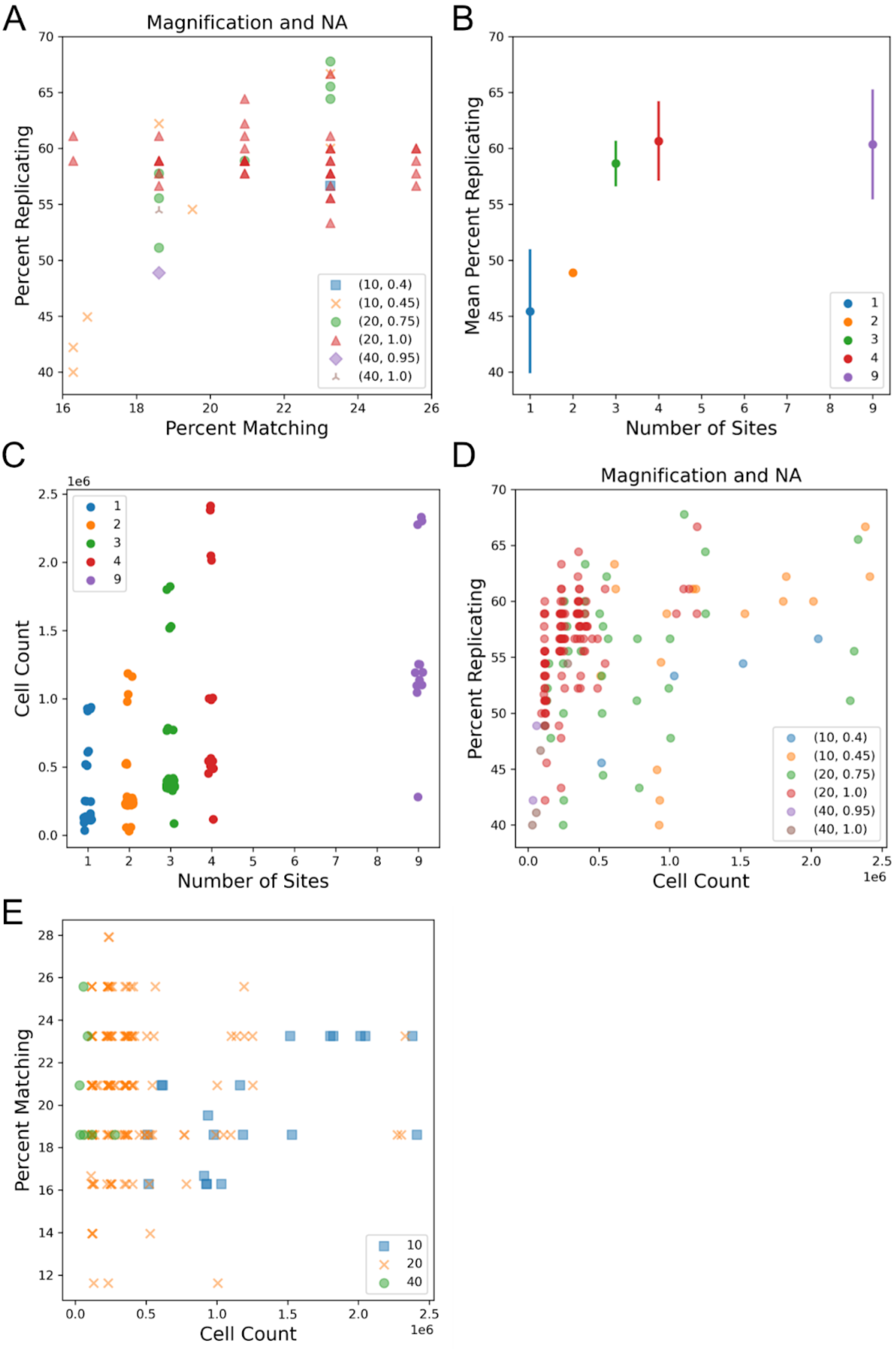
A
Magnification and NA
Percent Replicating
Percent Matching
(10, 0.4)
(10, 0.45)
(20, 0.75)
(20, 1.0)
(40, 0.95)
(40, 1.0)
B
Mean Percent Replicating
Number of Sites
1
2
3
4
9
C
Cell Count
1e6
Number of Sites
D
Magnification and NA
Percent Replicating
Cell Count
E
Percent Matching
Cell Count
10
20
40

**Supplementary figure 1**: Impact of magnification, number sites and cell count on image-based morphological profile technical quality.
(A) Comparison of profiles grouped based on magnification and numerical aperture (NA).
(B) Aggregated percent replicating score increases with the number of sites across all profiles.
(C) Cell count increases with the number of sites. Includes artificially site subsampled data.
(D) Profiles were grouped based on their magnification and aperture. An increase in cell count is associated with an increase in profile strength. Includes artificially site subsampled data. Cell count shown is sum across all wells.
(E) Comparison of percent matching versus cell count, grouped by magnification type. Includes artificially site subsampled data. Cell count shown is sum across all wells.

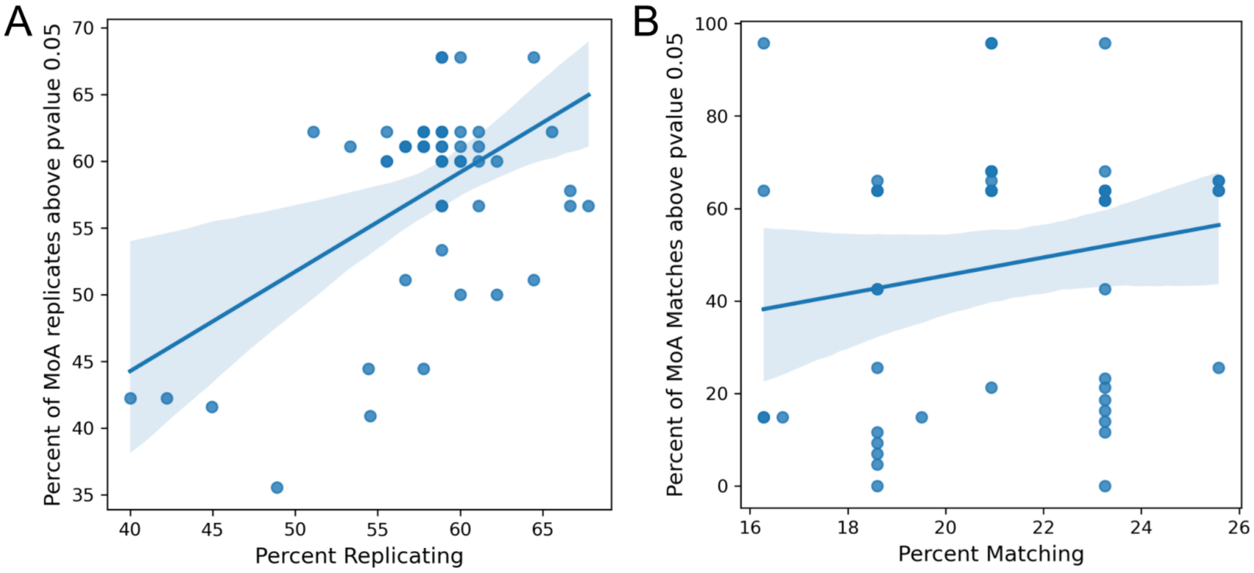


**Supplementary Figure 2**: Comparing mean average precision with percent replicating/matching for (A) replicate retrievability and (B) mechanism of action matching. The shaded area represents the 95% confidence interval for the linear regression between x and y.

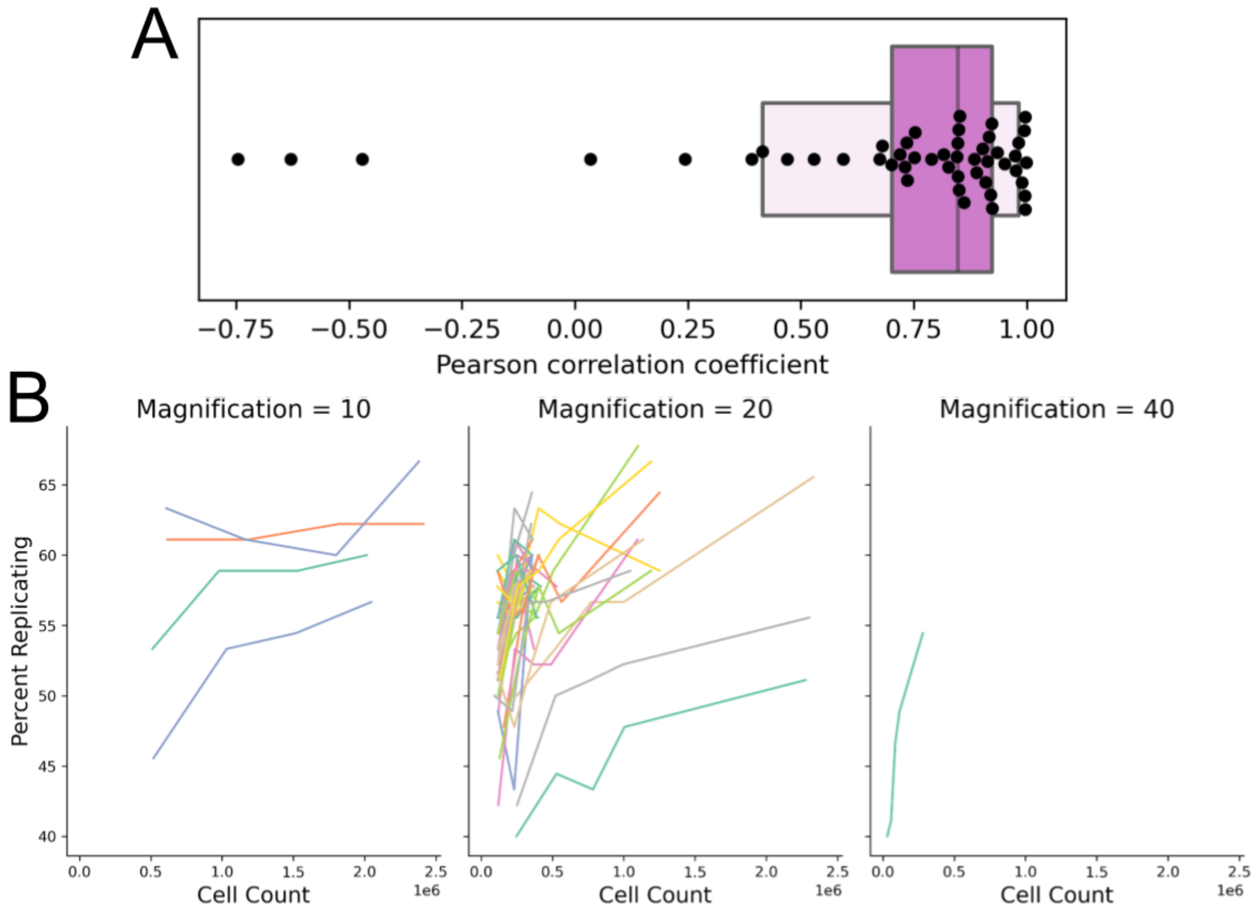


**Supplementary Figure 3**: Percent replicating increases with cell count.
(A) Pearson correlation coefficient values obtained from linear regression of percent replicating and cell count. Linear regressions were performed per-plate between profiles calculated on all sites and profiles calculated from subsamples. Only plates with at least 3 sites per well were used in this analysis. Each point represents the Pearson correlation coefficient of a linear regression for a given plate and its subsamples. In purple is the boxen plot (aka letter-value-plot) for this distribution. The central box represents the 25th to 75th percentile as in a standard box plot.
(B) Relationship between cell count and percent replicating for individual profiles + subsamples summarized in panel A. Plots are faceted based on magnification. Cell count shown is sum across all wells.

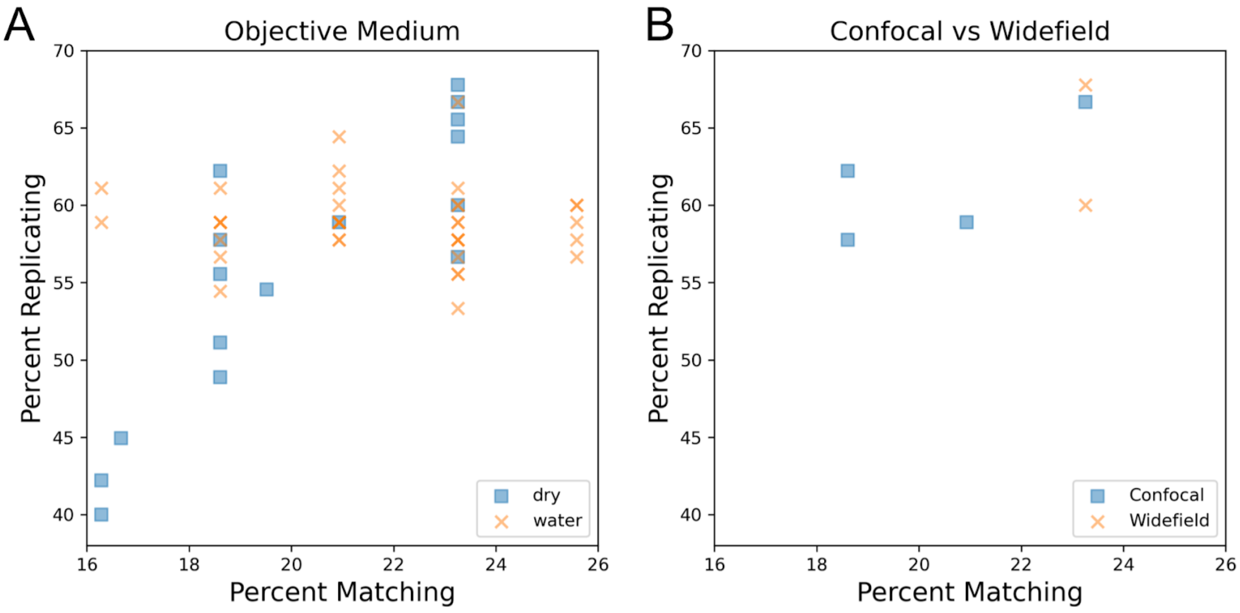


**Supplementary figure 4**: Exploration of the impact objective medium and imaging modality type has on image-based morphological profiles.

(A) Objective medium has minimal impact on profile strength.

(B) Comparison of additional confocal vs widefield profiles.

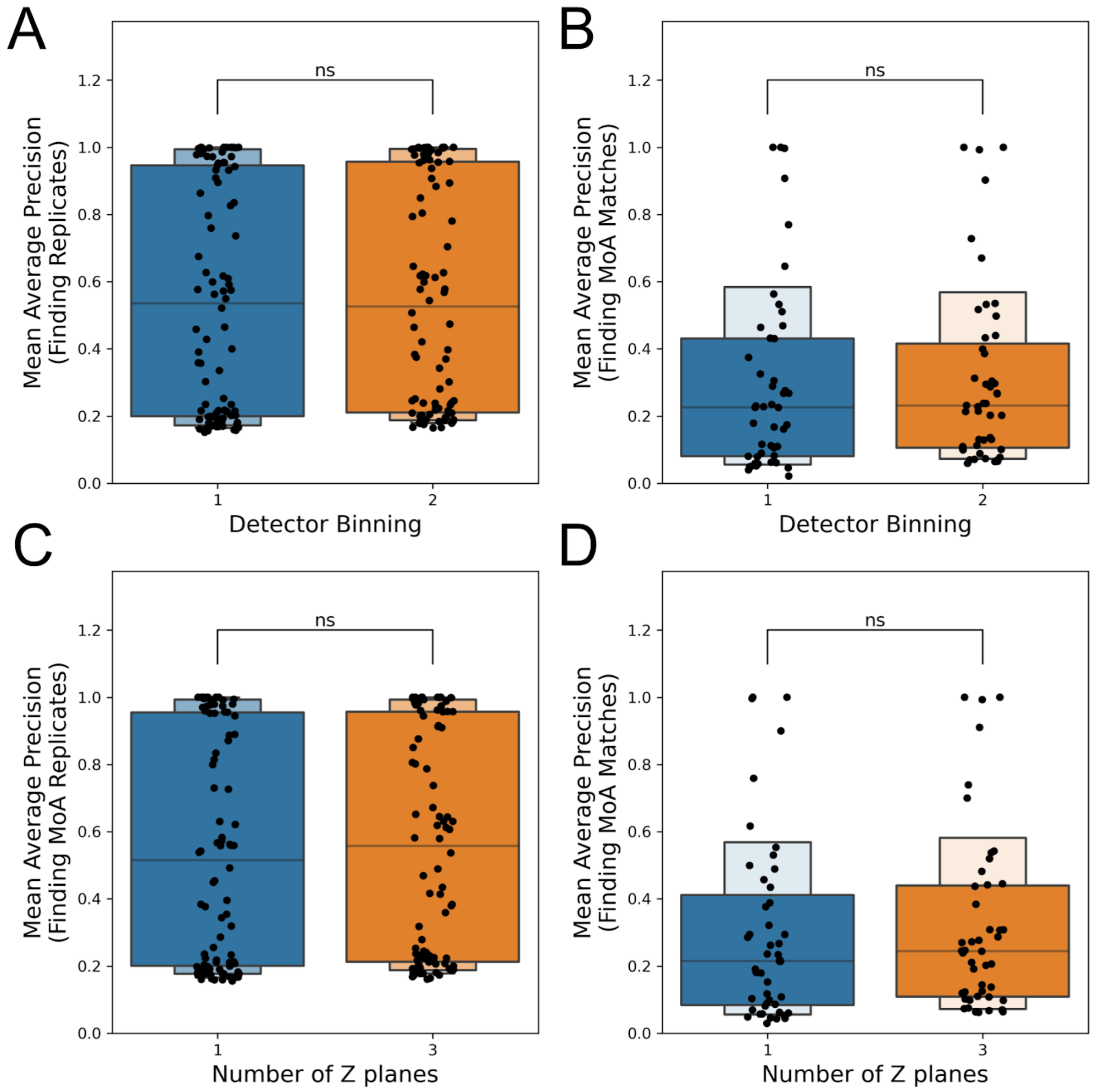


**Supplementary Figure 5**: mAP metrics show no significant difference between (A) detector binning or (B) number of z planes acquired.

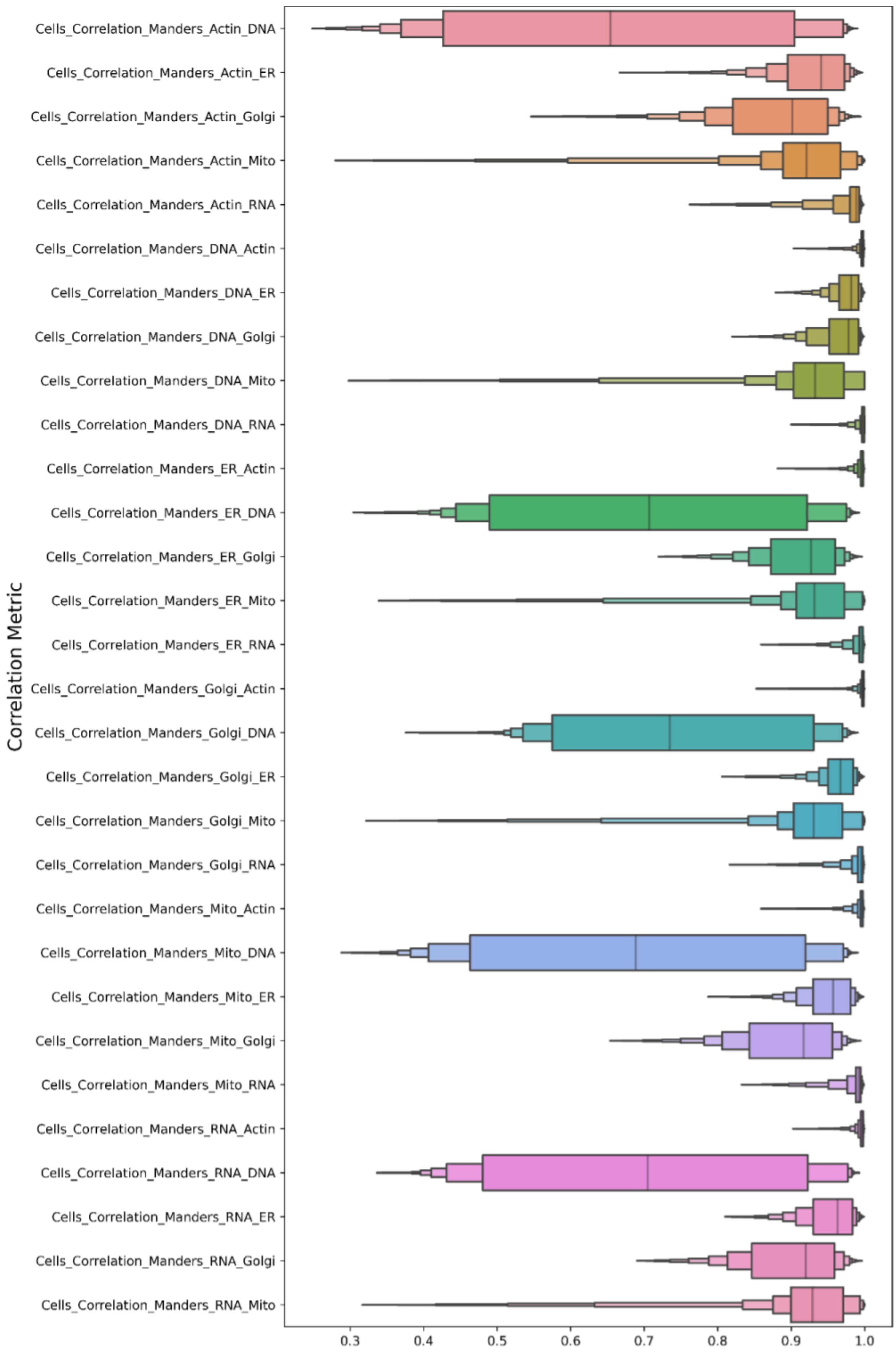
Correlation Metric
Cells_Correlation_Manders_Actin_DNA
Cells_Correlation_Manders_Actin_ER
Cells_Correlation_Manders_Actin_Golgi
Cells_Correlation_Manders_Actin_Mito
Cells_Correlation_Manders_Actin_RNA
Cells_Correlation_Manders_DNA_Actin
Cells_Correlation_Manders_DNA_ER
Cells_Correlation_Manders_DNA_Golgi
Cells_Correlation_Manders_DNA_Mito
Cells_Correlation_Manders_DNA_RNA
Cells_Correlation_Manders_ER_Actin
Cells_Correlation_Manders_ER_DNA
Cells_Correlation_Manders_ER_Golgi
Cells_Correlation_Manders_ER_Mito
Cells_Correlation_Manders_ER_RNA
Cells_Correlation_Manders_Golgi_Actin
Cells_Correlation_Manders_Golgi_DNA
Cells_Correlation_Manders_Golgi_ER
Cells_Correlation_Manders_Golgi_Mito
Cells_Correlation_Manders_Golgi_RNA
Cells_Correlation_Manders_Mito_Actin
Cells_Correlation_Manders_Mito_DNA
Cells_Correlation_Manders_Mito_ER
Cells_Correlation_Manders_Mito_Golgi
Cells_Correlation_Manders_Mito_RNA
Cells_Correlation_Manders_RNA_Actin
Cells_Correlation_Manders_RNA_DNA
Cells_Correlation_Manders_RNA_ER
Cells_Correlation_Manders_RNA_Golgi
Cells_Correlation_Manders_RNA_Mito
0.3
0.4
0.5
0.6
0.7
0.8
0.9
1.0

**Supplementary figure 6**: Unnormalized Manders correlation features within cell objects for all wells of profiles that split AGP into actin and Golgi + plasma membrane channels. High correlation is observed between actin and Golgi (especially above, Golgi_Actin, indicating that actin bleeds through into the Golgi channel), which are spectrally difficult to dissect.

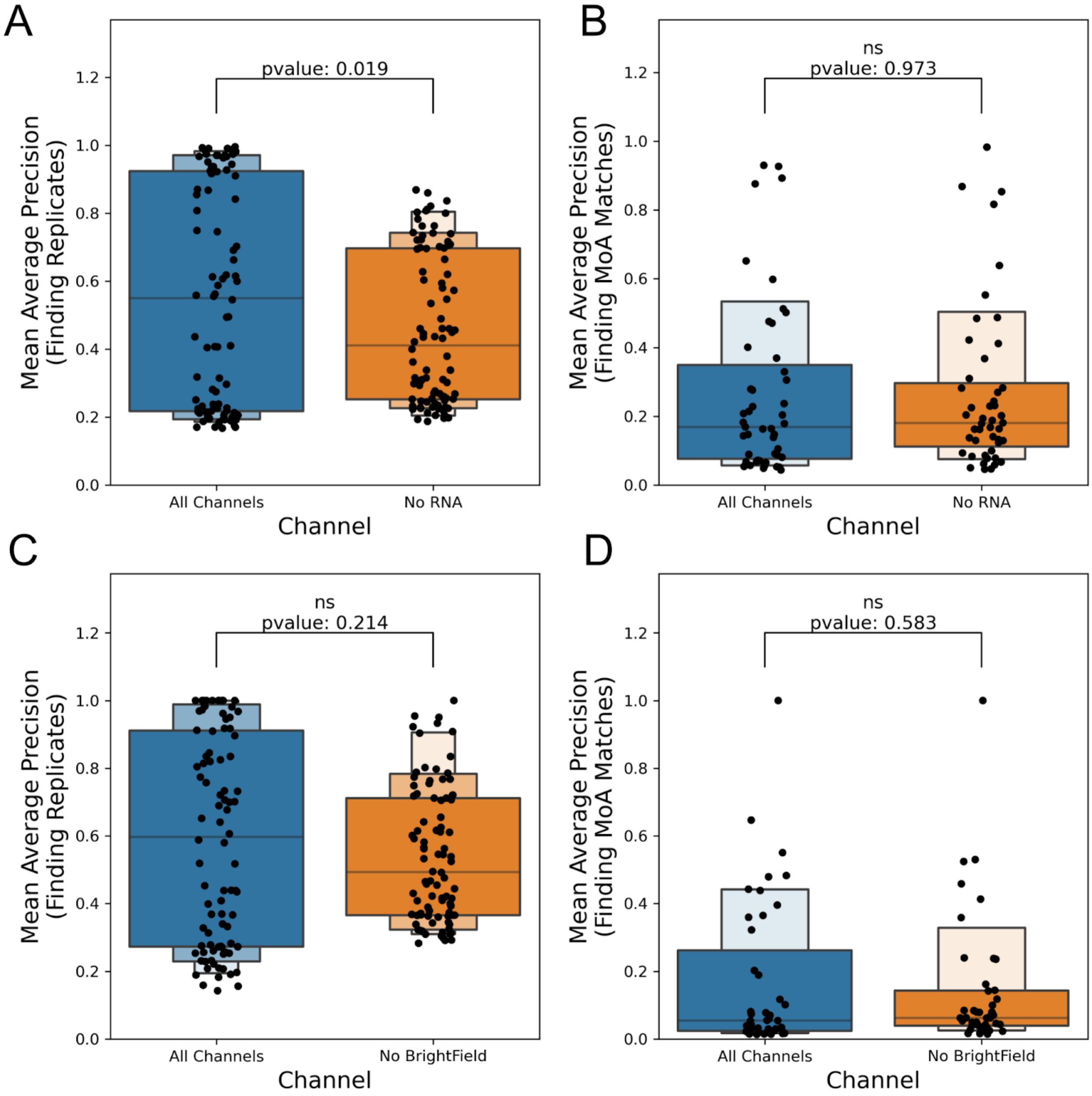


**Supplementary figure 7**: Alternative metric mAP used to compare profiles that have either (AB) RNA or (CD) brightfield channels dropped. Channels were dropped from negative control normalized profiles prior to feature selection. mAP was calculated between compound replicates (AC) and compounds with matching MoA (BD). All profiles were mean aggregated by compound for replicates or MoA for matching.

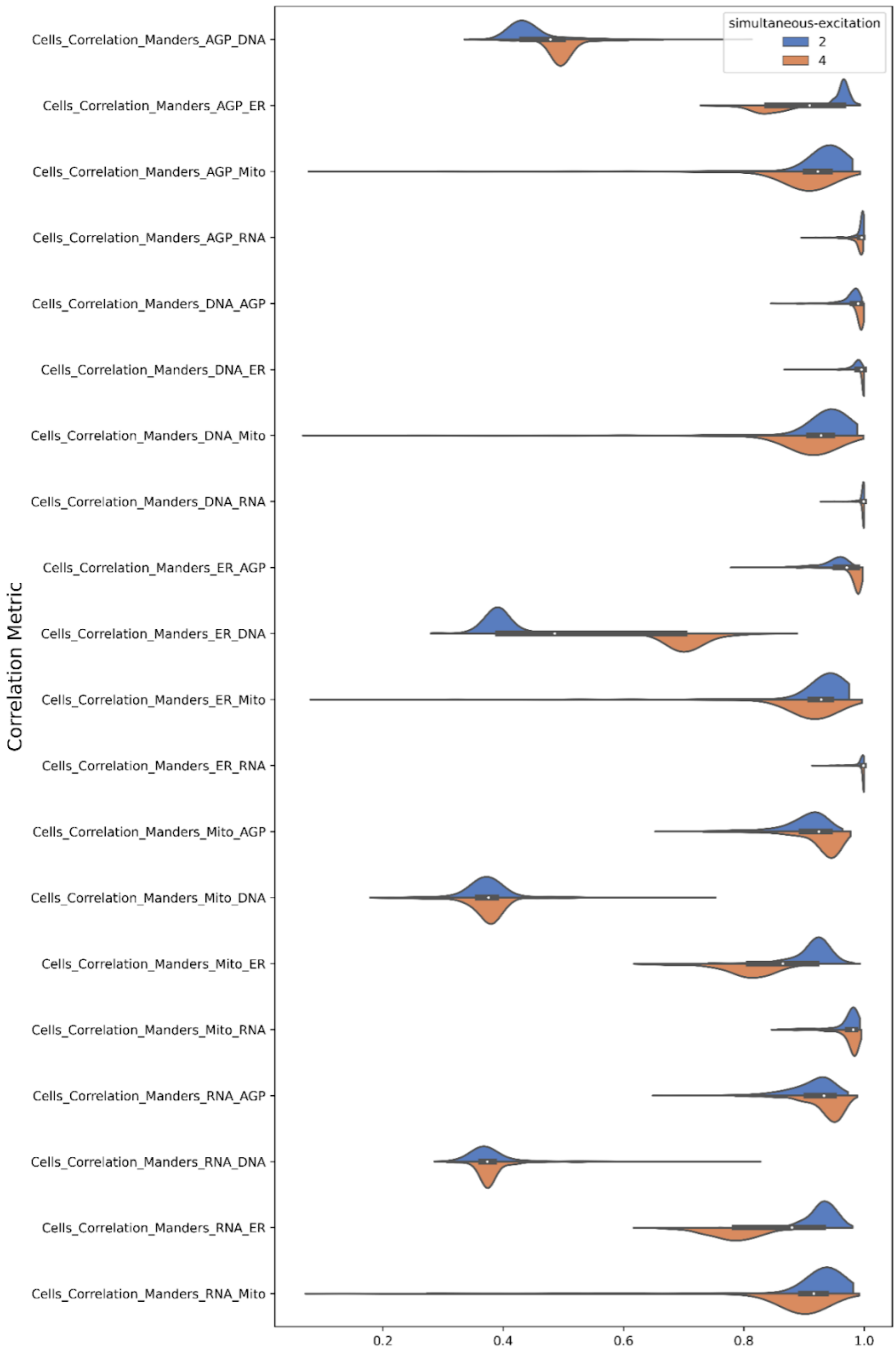
simultaneous-excitation
2
4
Correlation Metric
Cells_Correlation_Manders_AGP_DNA
Cells_Correlation_Manders_AGP_ER
Cells_Correlation_Manders_AGP_Mito
Cells_Correlation_Manders_AGP_RNA
Cells_Correlation_Manders_DNA_AGP
Cells_Correlation_Manders_DNA_ER
Cells_Correlation_Manders_DNA_Mito
Cells_Correlation_Manders_DNA_RNA
Cells_Correlation_Manders_ER_AGP
Cells_Correlation_Manders_ER_DNA
Cells_Correlation_Manders_ER_Mito
Cells_Correlation_Manders_ER_RNA
Cells_Correlation_Manders_Mito_AGP
Cells_Correlation_Manders_Mito_DNA
Cells_Correlation_Manders_Mito_ER
Cells_Correlation_Manders_Mito_RNA
Cells_Correlation_Manders_RNA_AGP
Cells_Correlation_Manders_RNA_DNA
Cells_Correlation_Manders_RNA_ER
Cells_Correlation_Manders_RNA_Mito
0.2
0.4
0.6
0.8
1.0

**Supplementary figure 8**: Unnormalized Manders correlation features within cell objects for 2 or 4x simultaneous excitation profiles. One plate for each condition is presented above. We observe an increase in DNA fluorescence bleedthrough into the ER channel (indicated by the ER_DNA feature above).

A

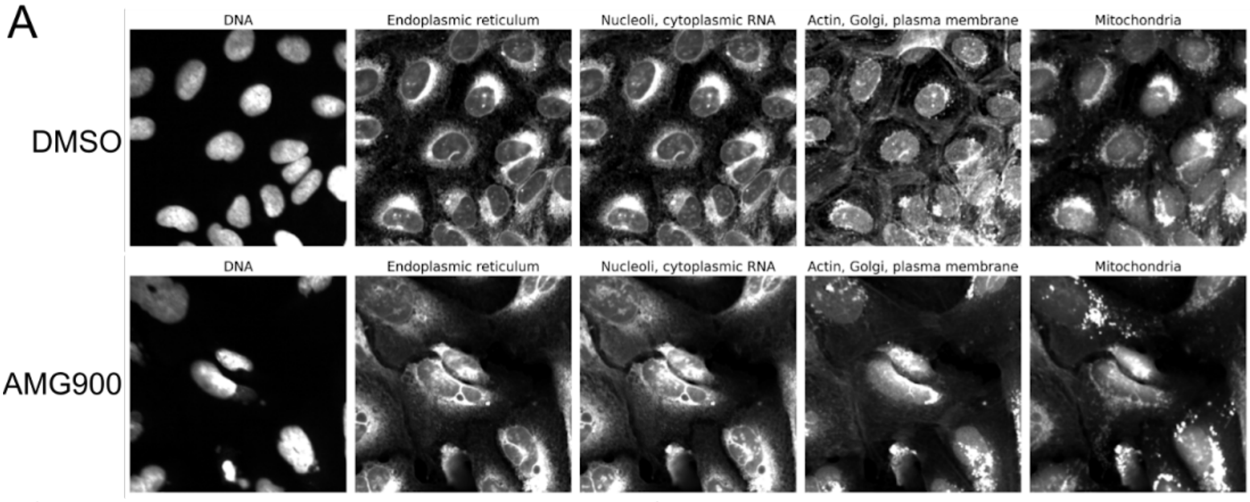


B

ImageXpress Micro Confocal

| | Feature | t-test |
|---|---|---|
| 1 | Cells_Intensity_IntegratedIntensityEdge_ER | 53.67 |
| 2 | Nuclei_AreaShape_Zernike_6_2 | 47.45 |
| 3 | Nuclei_AreaShape_Zernike_8_8 | 45.91 |
| 4 | Nuclei_Granularity_7_DNA | 36.33 |
| 5 | Cytoplasm_Intensity_IntegratedIntensityEdge_ER | 36.29 |
| 6 | Cytoplasm_Granularity_9_DNA | 35.63 |
| 7 | Cells_Granularity_9_DNA | 31.06 |
| 8 | Cells_Neighbors_FirstClosestObjectNumber_Adjacent | 31.03 |
| 9 | Cytoplasm_Granularity_10_DNA | 30.98 |
| 10 | Nuclei_AreaShape_BoundingBoxArea | 30.87 |

C

Eclipse Ti2

| | Feature | t-test |
|---|---|---|
| 1 | Cells_Granularity_4_DNA | 89.91 |
| 2 | Cells_Texture_DifferenceVariance_DNA_20_03_256 | 61.73 |
| 3 | Nuclei_AreaShape_Zernike_2_2 | 61.29 |
| 4 | Nuclei_Neighbors_FirstClosestDistance_2 | 54.33 |
| 5 | Cytoplasm_Granularity_4_DNA | 54.26 |
| 6 | Cells_RadialDistribution_FracAtD_DNA_3of4 | 53.46 |
| 7 | Cells_Intensity_UpperQuartileIntensity_DNA | 48.65 |
| 8 | Nuclei_Intensity_MassDisplacement_DNA | 47.58 |
| 9 | Nuclei_AreaShape_Zernike_0_0 | 45.51 |
| 10 | Nuclei_Granularity_1_ER | 45.14 |

D

Opera Phenix Plus

| | Feature | t-test |
|---|---|---|
| 1 | Cytoplasm_Granularity_8_DNA | 60.88 |
| 2 | Cells_Granularity_6_DNA | 42.11 |
| 3 | Cytoplasm_Granularity_14_DNA | 40.47 |
| 4 | Nuclei_AreaShape_Compactness | 38.4 |
| 5 | Cells_Texture_Correlation_DNA_20_03_256 | 37.6 |
| 6 | Nuclei_AreaShape_Zernike_8_8 | 36.55 |
| 7 | Nuclei_AreaShape_Solidity | 36.37 |
| 8 | Nuclei_AreaShape_Zernike_2_2 | 35.36 |
| 9 | Nuclei_AreaShape_Zernike_7_7 | 33.94 |
| 10 | Cytoplasm_Granularity_9_DNA | 33.26 |

E

Yokogawa CQ1

| | Feature | t-test |
|---|---|---|
| 1 | Nuclei_Intensity_MassDisplacement_DNA | 82.79 |
| 2 | Cells_Granularity_6_DNA | 74.45 |
| 3 | Nuclei_Granularity_9_DNA | 53.74 |
| 4 | Cytoplasm_Granularity_7_DNA | 50.66 |
| 5 | Cytoplasm_Granularity_2_RNA | 45.79 |
| 6 | Nuclei_Neighbors_AngleBetweenNeighbors_2 | 44.04 |
| 7 | Cells_Texture_Correlation_RNA_10_01_256 | 43.59 |
| 8 | Nuclei_AreaShape_FormFactor | 42.47 |
| 9 | Cytoplasm_Granularity_6_DNA | 42.35 |
| 10 | Nuclei_Texture_Correlation_DNA_20_01_256 | 42.24 |

F

Yokogawa CV8000

| | Feature | t-test |
|---|---|---|
| 1 | Cells_Texture_Correlation_RNA_20_03_256 | 90.68 |
| 2 | Cells_RadialDistribution_FracAtD_BrightField_4of4 | 60.0 |
| 3 | Cells_RadialDistribution_FracAtD_BrightField_3of4 | 53.87 |
| 4 | Cells_Texture_Correlation_Mito_20_00_256 | 53.39 |
| 5 | Cells_Neighbors_FirstClosestDistance_Adjacent | 52.18 |
| 6 | Cytoplasm_Granularity_7_DNA | 50.78 |
| 7 | Nuclei_AreaShape_Zernike_2_0 | 48.38 |
| 8 | Nuclei_Granularity_7_DNA | 46.3 |
| 9 | Cytoplasm_Texture_InfoMeas2_ER_5_00_256 | 42.67 |
| 10 | Cells_AreaShape_Extent | 42.55 |

**Supplementary Figure 9**: Marker selection between DMSO and AMG900 treated cells
(A) Representative images of U2OS cells treated with negative control DMSO and the Aurora kinase inhibitor AMG900 (3 μM)
(B-F) Marker selection was used to examine which features differentiate the most between DMSO and AMG900 treatments for (B) Molecular Devices ImageXpress Micro Confocal, (C) Nikon Eclipse Ti2, (D) Revvity Opera Phenix Plus, (E) Yokogawa CQ1, and (F) Yokogawa CV8000 imaging systems.

A

### Molecular Devices ImageXpress Micro Confocal

Least sensitive features

| | Feature | pvalue |
|---|---|---|
| 1 | Cytoplasm_AreaShape_Eccentricity | 0.330 |
| 2 | Cells_Neighbors_SecondClosestDistance | 0.217 |
| 3 | Cytoplasm_AreaShape_MeanRadius | 0.197 |
| 4 | Cytoplasm_AreaShape_MedianRadius | 0.190 |
| 5 | Cytoplasm_AreaShape_MinorAxisLength | 0.165 |
| 6 | Cells_AreaShape_EquivalentDiameter | 0.163 |
| 7 | Cells_AreaShape_MinorAxisLength | 0.149 |
| 8 | Nuclei_Neighbors_SecondClosestDistance | 0.129 |
| 9 | Cytoplasm_AreaShape_MaximumRadius | 0.123 |
| 10 | Cells_AreaShape_FormFactor | 0.123 |

Most sensitive features

| | Feature | pvalue |
|---|---|---|
| 1 | Cells_AreaShape_EulerNumber | 2.123e-12 |
| 2 | Nuclei_AreaShape_MedianRadius | 4.982e-09 |
| 3 | Cells_AreaShape_Extent | 7.404e-09 |
| 4 | Cells_Neighbors_NumberOfNeighbors | 8.722e-09 |
| 5 | Nuclei_AreaShape_MaximumRadius | 1.139e-08 |
| 6 | Nuclei_AreaShape_MinFeretDiameter | 1.742e-08 |
| 7 | Nuclei_AreaShape_MeanRadius | 2.652e-08 |
| 8 | Nuclei_AreaShape_MinorAxisLength | 4.009e-08 |
| 9 | Cytoplasm_Intensity_MassDisplacement | 2.293e-06 |
| 10 | Nuclei_AreaShape_EquivalentDiameter | 2.672e-06 |

B

### Nikon Eclipse Ti2

Least sensitive features

| | Feature | pvalue |
|---|---|---|
| 1 | Nuclei_AreaShape_Orientation | 0.422 |
| 2 | Cytoplasm_Granularity_16 | 0.382 |
| 3 | Cells_AreaShape_Area | 0.363 |
| 4 | Cells_Granularity_16 | 0.349 |
| 5 | Cells_AreaShape_EquivalentDiameter | 0.336 |
| 6 | Nuclei_Granularity_15 | 0.325 |
| 7 | Cytoplasm_AreaShape_BoundingBoxArea | 0.315 |
| 8 | Cells_AreaShape_BoundingBoxArea | 0.315 |
| 9 | Nuclei_Granularity_16 | 0.311 |
| 10 | Nuclei_Neighbors_AngleBetweenNeighbors | 0.306 |

Most sensitive features

| | Feature | pvalue |
|---|---|---|
| 1 | Nuclei_AreaShape_MaximumRadius | 0.003 |
| 2 | Nuclei_AreaShape_Perimeter | 0.004 |
| 3 | Nuclei_AreaShape_EquivalentDiameter | 0.005 |
| 4 | Nuclei_AreaShape_MaxFeretDiameter | 0.005 |
| 5 | Nuclei_Intensity_MeanIntensityEdge | 0.006 |
| 6 | Nuclei_AreaShape_Area | 0.006 |
| 7 | Cytoplasm_Intensity_MeanIntensityEdge | 0.006 |
| 8 | Nuclei_Intensity_IntegratedIntensityEdge | 0.006 |
| 9 | Nuclei_AreaShape_MeanRadius | 0.006 |
| 10 | Nuclei_AreaShape_MinorAxisLength | 0.006 |

C

### Revvity Opera Phenix Plus

Least sensitive features

| | Feature | pvalue |
|---|---|---|
| 1 | Nuclei_AreaShape_Orientation | 0.372 |
| 2 | Cells_AreaShape_Orientation | 0.366 |
| 3 | Cells_AreaShape_BoundingBoxMaximum | 0.355 |
| 4 | Cytoplasm_AreaShape_BoundingBoxMaximum | 0.355 |
| 5 | Nuclei_AreaShape_BoundingBoxMaximum | 0.349 |
| 6 | Nuclei_AreaShape_Center | 0.342 |
| 7 | Cells_AreaShape_Center | 0.340 |
| 8 | Cytoplasm_AreaShape_Center | 0.336 |
| 9 | Cells_AreaShape_BoundingBoxMinimum | 0.324 |
| 10 | Cytoplasm_AreaShape_BoundingBoxMinimum | 0.324 |

Most sensitive features

| | Feature | pvalue |
|---|---|---|
| 1 | Nuclei_Intensity_IntegratedIntensityEdge | 0.009 |
| 2 | Cytoplasm_Intensity_MaxIntensity | 0.009 |
| 3 | Nuclei_Intensity_MaxIntensityEdge | 0.010 |
| 4 | Nuclei_Texture_SumEntropy | 0.010 |
| 5 | Nuclei_Texture_Entropy | 0.010 |
| 6 | Nuclei_Intensity_IntegratedIntensity | 0.010 |
| 7 | Nuclei_Intensity_MedianIntensity | 0.010 |
| 8 | Nuclei_Intensity_MeanIntensity | 0.010 |
| 9 | Nuclei_Intensity_MaxIntensity | 0.011 |
| 10 | Nuclei_Texture_SumAverage | 0.011 |

D

### Yokogawa CV8000

Least sensitive features

| | Feature | pvalue |
|---|---|---|
| 1 | Nuclei_AreaShape_Orientation | 0.307 |
| 2 | Cytoplasm_AreaShape_Orientation | 0.277 |
| 3 | Cells_Neighbors_FirstClosestObjectNumber | 0.264 |
| 4 | Cells_Neighbors_SecondClosestObjectNumber | 0.262 |
| 5 | Nuclei_Neighbors_FirstClosestObjectNumber | 0.258 |
| 6 | Cells_AreaShape_Orientation | 0.256 |
| 7 | Nuclei_Neighbors_SecondClosestObjectNumber | 0.254 |
| 8 | Cells_Neighbors_FirstClosestDistance | 0.198 |
| 9 | Cells_AreaShape_BoundingBoxMaximum | 0.184 |
| 10 | Cytoplasm_AreaShape_BoundingBoxMaximum | 0.184 |

Most sensitive features

| | Feature | pvalue |
|---|---|---|
| 1 | Nuclei_Intensity_StdIntensityEdge | 2.998e-07 |
| 2 | Cytoplasm_Texture_Variance | 4.907e-06 |
| 3 | Cytoplasm_AreaShape_Compactness | 6.100e-06 |
| 4 | Cytoplasm_Texture_SumVariance | 6.671e-06 |
| 5 | Nuclei_Intensity_StdIntensity | 6.887e-06 |
| 6 | Cytoplasm_Intensity_MaxIntensity | 1.989e-05 |
| 7 | Cytoplasm_Texture_SumEntropy | 2.847e-05 |
| 8 | Cytoplasm_Texture_Contrast | 3.164e-05 |
| 9 | Nuclei_Intensity_LowerQuartileIntensity | 3.459e-05 |
| 10 | Nuclei_Intensity_MeanIntensityEdge | 7.672e-05 |

E

### Yokogawa CV8000 - Same simultaneous excitation

Least sensitive features

| | Feature | pvalue |
|---|---|---|
| 1 | Cells_Neighbors_FirstClosestObjectNumber | 0.501 |
| 2 | Nuclei_Neighbors_FirstClosestObjectNumber | 0.496 |
| 3 | Cells_Neighbors_SecondClosestObjectNumber | 0.491 |
| 4 | Nuclei_Neighbors_SecondClosestObjectNumber | 0.479 |
| 5 | Cells_Neighbors_SecondClosestDistance | 0.333 |
| 6 | Nuclei_Neighbors_SecondClosestDistance | 0.331 |
| 7 | Nuclei_AreaShape_Center | 0.321 |
| 8 | Nuclei_AreaShape_BoundingBoxMinimum | 0.315 |
| 9 | Cells_AreaShape_Center | 0.315 |
| 10 | Cells_Neighbors_FirstClosestDistance | 0.301 |

Most sensitive features

| | Feature | pvalue |
|---|---|---|
| 1 | Cells_Neighbors_PercentTouching | 1.216e-16 |
| 2 | Cytoplasm_Granularity_3 | 4.022e-12 |
| 3 | Nuclei_Intensity_MinIntensity | 2.961e-11 |
| 4 | Cytoplasm_AreaShape_Compactness | 9.063e-11 |
| 5 | Nuclei_AreaShape_MeanRadius | 2.405e-10 |
| 6 | Nuclei_AreaShape_MaximumRadius | 3.886e-10 |
| 7 | Cytoplasm_Intensity_IntegratedIntensity | 9.128e-10 |
| 8 | Cells_AreaShape_Compactness | 1.586e-09 |
| 9 | Nuclei_AreaShape_Compactness | 1.586e-09 |
| 10 | Cells_AreaShape_MajorAxisLength | 2.583e-09 |

**Supplementary Figure 10**: Exploring which features are least and most sensitive for each imaging system; see methods for more information. The top (most sensitive) and bottom (least sensitive) 10 p-value features are shown for:
(A) Molecular Devices ImageXpress Micro Confocal
(B) Nikon Eclipse Ti2
(C) Revvity Opera Phenix Plus
(D) Yokogawa CV8000
(E) Yokogawa CV8000, only comparing within profiles with the same simultaneous excitation settings. Since profiles captured with 4 simultaneous excitations will contain large levels of bleedthrough that profiles captured with 2 simultaneous excitations will not have (see Figure 4BC), dropping comparisons between pairs of plates where one plate had such bleedthrough and one did not leaves only 2 intensity features in the top 10 most sensitive across settings, rather than 5 without such correction.

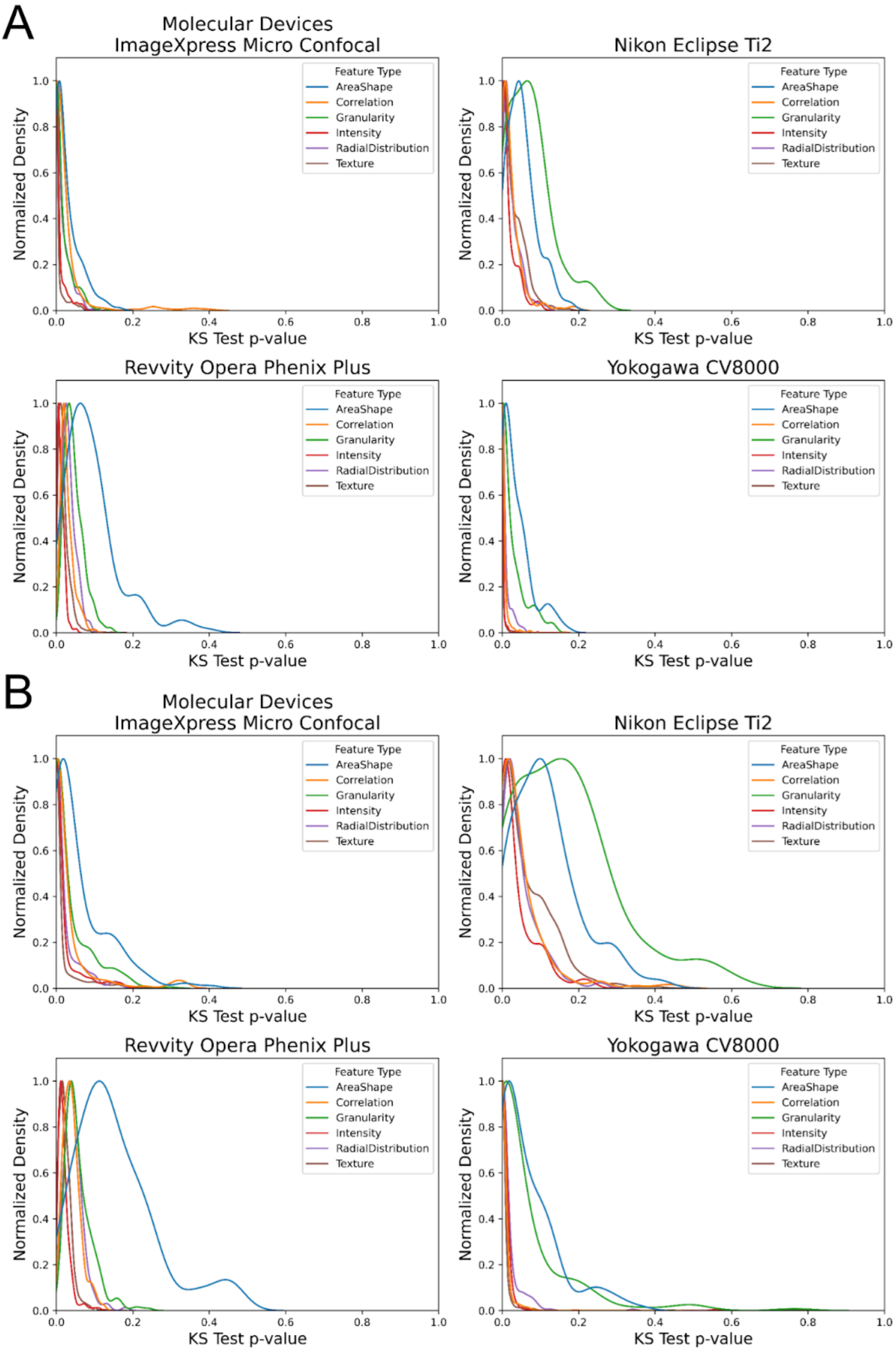

A
Molecular Devices
ImageXpress Micro Confocal
Nikon Eclipse Ti2
Revvity Opera Phenix Plus
Yokogawa CV8000
B
Molecular Devices
ImageXpress Micro Confocal
Nikon Eclipse Ti2
Revvity Opera Phenix Plus
Yokogawa CV8000
Feature Type
AreaShape
Correlation
Granularity
Intensity
RadialDistribution
Texture
Normalized Density
KS Test p-value
0.0
0.2
0.4
0.6
0.8
1.0

**Supplementary Figure 11**: A two-sample KS test was performed between all features for (A) all possible combinations of plates within a vendor or (B) combinations of plates that were acquired at the same pixel size (i.e. the same magnification and detector binning settings); see Methods. Density is normalized for each feature group so that the maximum value is 1.
